# Germline-mediated immunoediting sculpts breast cancer subtypes and metastatic proclivity

**DOI:** 10.1101/2023.03.15.532870

**Authors:** Kathleen E. Houlahan, Aziz Khan, Noah F Greenwald, Robert B. West, Michael Angelo, Christina Curtis

**Affiliations:** Stanford Cancer Institute, Stanford University School of Medicine, Stanford, CA, USA; Cancer Biology Program, Stanford University School of Medicine, Stanford, CA, USA; Department of Pathology, Stanford University School of Medicine, Stanford, CA, USA; Department of Medicine (Oncology), Stanford University School of Medicine, Stanford, CA, USA; Department of Genetics, Stanford University School of Medicine, Stanford, CA, USA; Department of Biomedical Data Science, Stanford University School of Medicine, Stanford, CA, USA; Chan Zuckerberg Biohub, San Francisco, CA, USA

## Abstract

Cancer represents a broad spectrum of molecularly and morphologically diverse diseases. Individuals with the same clinical diagnosis can have tumors with drastically different molecular profiles and clinical response to treatment. It remains unclear when these differences arise during disease course and why some tumors are addicted to one oncogenic pathway over another. Somatic genomic aberrations occur within the context of an individual’s germline genome, which can vary across millions of polymorphic sites. An open question is whether germline differences influence somatic tumor evolution. Interrogating 3,855 breast cancer lesions, spanning pre-invasive to metastatic disease, we demonstrate that germline variants in highly expressed and amplified genes influence somatic evolution by modulating immunoediting at early stages of tumor development. Specifically, we show that the burden of germline-derived epitopes in recurrently amplified genes selects against somatic gene amplification in breast cancer. For example, individuals with a high burden of germline-derived epitopes in *ERBB2,* encoding human epidermal growth factor receptor 2 (HER2), are significantly less likely to develop HER2-positive breast cancer compared to other subtypes. The same holds true for recurrent amplicons that define four subgroups of ER-positive breast cancers at high risk of distant relapse. High epitope burden in these recurrently amplified regions is associated with decreased likelihood of developing high risk ER-positive cancer. Tumors that overcome such immune-mediated negative selection are more aggressive and demonstrate an “immune cold” phenotype. These data show the germline genome plays a previously unappreciated role in dictating somatic evolution. Exploiting germline-mediated immunoediting may inform the development of biomarkers that refine risk stratification within breast cancer subtypes.

## Introduction

Malignancy is defined by a set of abnormal biological capacities, termed the hallmarks of cancer (*1*). Decades of histopathologic assessment and molecular profiling of human tumors have demonstrated there are multiple ways cells can acquire each hallmark (*2, 3*). As a result, tumors with the same clinical characteristics can vary dramatically across individuals and these distinct molecular vulnerabilities can have important prognostic and therapeutic implications. It is unclear when these differences originate.

Oncogenic aberrations are acquired within the context of germline genomes which differ across individuals at millions of polymorphic sites (*4*), but the role of germline variants in somatic evolution remains poorly understood. The most compelling example is that deleterious germline variants in BRCA1 and, to a lesser extent, BRCA2 are preferentially associated with the development of triple negative breast cancer (TNBC) (*5*), implying germline variants sculpt specific subtypes of disease (*6, 7*). The mechanistic basis for this preference is incompletely characterized. Additionally, germline variants that upregulate the mTOR pathway are associated with further deregulation of mTOR *via* somatic PTEN loss-of-function (*8*). Moreover, pathogenic germline variants in cancer predisposition genes promote somatic bi-allelic inactivation in a lineage-dependent manner (*9*). In prostate cancer, germline variants can modulate genomic stability, tumor-specific DNA methylation and gene regulation at the transcriptional and translational levels (*10, 11*). Differences in breast cancer subtype frequencies across ancestral populations further suggest germline contributions (*12, 13*). These data point to an underappreciated role of the germline genome in somatic tumor evolution.

Various lines of evidence suggest that avoidance of the adaptive immune system is another strong determinant of which somatic mutations persist within a tumor (*14, 15*). It remains less clear how germline differences influence immunoediting. Levels of interferon signaling and cytotoxic T-cell infiltration are estimated to be 15-20% heritable (*16*). Generally, germline variants have not been considered a good source of immunogenic epitopes as cytotoxic response should be dampened by central tolerance. However, non-mutated immunogenic epitopes have been identified in genes such as *ERBB2/*HER2 in breast and ovarian cancer (*17*) and H4 histone in prostate cancer (*18*), amongst others (*19, 20*). Antigens with weak binding affinity for MHC receptors can escape central tolerance (*21*) and elicit an immune response (*22*). Tissue-restricted post-translational modifications can also circumvent central tolerance (*23*). Peripheral self-reactive T cells are present at similar frequencies to T cells specific to foreign antigens (*24*) but are held in an anergic state by regulatory T cells (Treg) (*25*). However, Treg depletion in healthy mouse models led to the natural occurrence of self-reactive CD4+ T cells (*26*). Further, there is mounting evidence for innate-like T-cell populations within mouse and tumor malignancies that have increased propensity for self-reactivity (*27*). Altogether, these data suggest that under specific circumstances and disruptions to immune homeostasis, a subset of T-cells may respond to germline-derived epitopes during tumorigenesis.

Building on these observations, we sought to investigate whether germline variants sculpt somatic evolution by mediating immunoediting. Specifically, we hypothesize that the burden of germline-derived epitopes in recurrently amplified driver genes may select against gene amplification. This is because amplification of a gene with a high burden of germline-derived epitopes would increase epitope availability, likelihood of epitope presentation and immune-mediated cell death. Instead, immune pressures may select for amplification of an alternate driver gene with a lower germline-mediated epitope burden. We addressed this question in breast cancer for three reasons. First, the well-characterized link between BRCA1 and TNBC susceptibility (*5*), along with high heritability estimates (∼31%) (*28*), suggests the germline genome plays a role in shaping breast cancer evolution. Second, breast cancer is one of the most extensively sequenced cancer types with sizeable cohorts spanning the full continuum of disease, from pre-invasive lesions to primary tumors and metastatic disease (*2, 3, 29–31*). Finally, oncogenic amplifications define five prognostic breast cancer subtypes, including the *ERBB2/*HER2-positive (HER2+) subgroup and four estrogen receptor (ER) positive, HER2-negative (ER+/HER2-) genomic subgroups (*32, 33*) which are established early, evidenced by their identification in premalignant ductal carcinoma *in situ* (DCIS) (*30*). Thus, breast cancer provides an optimal proof-of-concept for studying this phenomenon.

We leveraged paired tumor and normal sequencing data from 1,087 primary (*3, 31*) and 702 metastatic (*29*) breast cancer patients as well as somatic genomic profiles from 341 patients with DCIS (*30*) and evaluated the relationship between germline-derived epitope burden (henceforth referred to as GEB) and subtype commitment, defined by the acquisition of focal oncogenic amplifications. We found that high GEB in subtype specific genes was consistently negatively associated with somatic gene amplification and, therefore, subtype commitment. Tumors that successfully overcome a high GEB were more aggressive and exhibited microenvironments depleted of lymphocytes, consistent with “immune cold” tumors. These data indicate that supposedly “benign” germline variants with little to no functional genic effect, may, in aggregate, sculpt breast cancer subtypes and disease aggression *via* immunoediting.

## Results

### Germline-derived epitope burden (GEB) selects against cognate oncogene amplification

Germline genomes, which vary from individual to individual across millions of polymorphic sites, serve as the substrate for the acquisition of somatic genomic alterations (*4*). We hypothesize that inherited variation in amplified oncogenes could result in varied immunoediting pressures across individuals. Specifically, if an individual harbors germline variants that produce epitopes that can be presented by their MHC class I, selection against genomic amplification of the germline epitopes may occur thus mitigating an anti-tumor immune response. A simplified example is illustrated in **Figure 1A**, where individual A does not have germline polymorphisms in gene X. Both copies of gene X in individual A match the reference genome exactly so we refer to this individual as wildtype (WT). When translated, protein X produces two peptides that can be presented by individual A’s MHC class I. Suppose during early stages of transformation, individual A acquires a somatic amplification such that there are three copies of gene X. In that case, this results in six available epitopes (2 unique epitopes x 3 gene copies = 6). Next, we consider individual B who has a homozygous polymorphism in gene X such that three peptides derived from protein X can be presented. If individual B gains three copies of gene X, nine epitopes are now produced. In contrast, individual C harbors a homozygous polymorphism that results in only one presentable epitope. In this case, amplification of gene X generates three epitopes. We assume the number of produced epitopes positively correlates with the likelihood of presentation. In this case, individual B would have the highest likelihood of presentation and the highest likelihood of triggering immune surveillance. In contrast, individual C has the lowest likelihood of presentation and immune surveillance. As a result, amplification of gene X would be more beneficial to individual C than individual B, who experiences an increased fitness cost when amplifying gene X.

**Figure 1.**
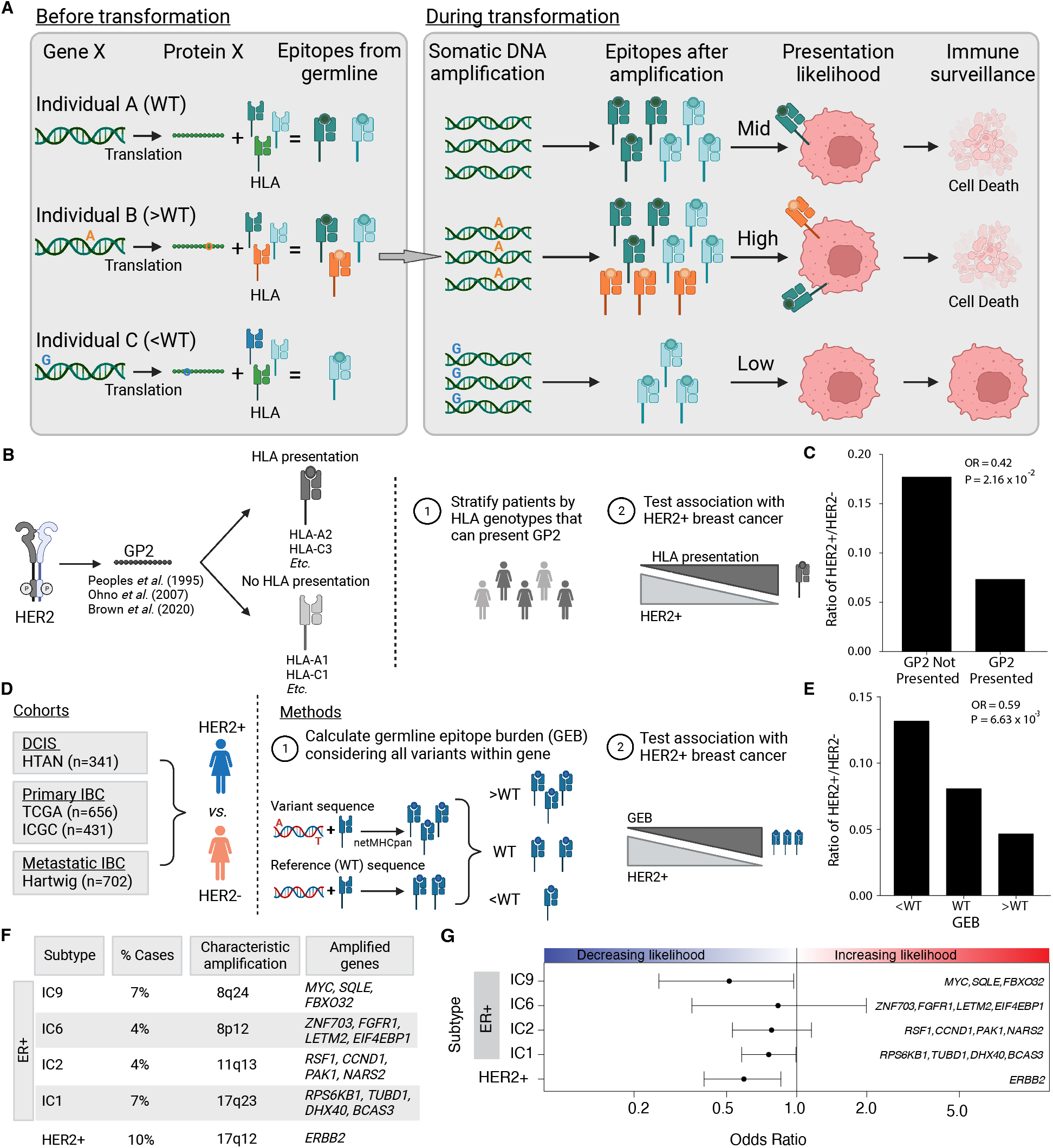
Germline-derived epitope burden in oncogenes selects against oncogene amplification. **A)** Schematic of germline-mediated immunoediting. Prior to transformation, differing germline genomes and HLA alleles result in differing numbers of epitopes derived from a gene of interest (*e.g.* Gene X). If during transformation, the tumor acquires additional copies of Gene X (*i.e.* somatic amplification), the number of epitopes increases further. As a result, individuals with a high burden of epitopes are more likely to be surveilled by the immune system triggering cell death. **B)** As proof of concept, GP2 is a well-characterized, naturally occurring (*i.e.,* non- somatically mutated) immunogenic peptide derived from HER2. Schematic overview of analysis framework to investigate if the ability to present GP2, *i.e.,* having MHC Class I alleles that bind GP2, is associated with HER2+ breast cancer. **C)** The ability to present GP2 is negatively associated with HER2+ breast cancer. Barplot shows the ratio of HER+ to HER2- in patients that have HLA alleles that can bind GP2 (GP2 presented) *vs* patients that do not (GP2 not presented). Odds ratio (OR) and p-value from logistic regression model correcting for first six genetic principal components. **D)** Schematic outlining methods to investigate germline-mediated immunoediting. Using four independent cohorts representing multiple stages of breast cancer, pre- invasive, primary invasive and metastatic invasive breast cancer, we investigated if the GEB in a gene of interest was associated with the likelihood of acquiring a somatic amplification of the gene using HER2 as a representative example. **E)** GEB in *ERBB2* is negatively associated with HER2+ breast cancer. Barplot shows the ratio of HER2+ to HER2- patients with low, medium or high GEB. Odds ratio (OR) and p-value from logistic regression model correcting for the first six genetic principal components. **F)** Beyond *ERRB2*, we investigated amplicons that characterize four ER+/HER2- high risk of relapse subtypes (IC1, IC2, IC6 and IC9), where the percent of breast cancer cases they represent and the corresponding chromosome region and core genes is denoted for each subtype. **G)** GEB in recurrently amplified genes is negatively associated with gene amplification. Scatterplot shows odds ratio (x-axis) and 95% confidence intervals from logistic regression model correcting for the first six genetic principal components and somatic mutation burden. Covariates in the top panel indicate the direction of the effect, namely whether GEB is associated with increased or decreased likelihood of each subtype.

If this scenario occurs in human tumors, evidence of germline epitope-mediated negative selection of oncogenes should be detectable in retrospective cancer sequencing data. As proof of concept, we first considered *ERBB2* as gene X. We identified, from the literature, a non-somatically mutated immunogenic peptide derived from HER2 that has been extensively characterized (*17, 34, 35*). GP2 is a nine amino acid immunogenic peptide derived from the transmembrane domain of HER2 (aa 654-662; IISAVVGIL). GP2 has been repeatedly shown to elicit an immune response in various cancer settings, i.e., breast, ovarian and hepatocellular cancer (*17, 35*). A multi-center phase II clinical trial (NCT00524277) found that HER2+ breast cancer patients treated with the GP2 vaccine (nvaccine = 48) experienced no recurrences in contrast to the untreated control group (ncontrol = 50; 100% vs 87.2%; P = 0.052) (*34*). Based on these promising fundings, a randomized Phase II trial (NCT05232916) is underway. Given its established immunogenicity, we evaluated whether the naturally occurring GP2 influences development of HER2+ breast cancer. Specifically, we asked if ability to present GP2 (*i.e.,* having HLA alleles that bind endogenous GP2 peptide) was associated with HER2+ breast cancer in individuals of European descent from two primary breast cancer cohorts: the International Cancer Genome Consortium (ICGC; n = 431) (*31*) and The Cancer Genome Atlas (TCGA; n = 656) (**Figure 1B**) (*3*). Indeed, we found that individuals that had HLA alleles that could bind GP2 were significantly less likely to develop HER2+ breast cancer in both ICGC (**Figure 1C)** and TCGA (meta-analysis: OR = 0.60; P = 0.087; **Supplementary Figure 1A**). These data illustrate that germline epitopes play a role in dictating breast cancer subtypes motivating more comprehensive interrogation of the phenomenon.

Building on this proof-of-concept, we investigated if germline-mediated negative selection extends beyond GP2 and considered germline-derived epitopes throughout *ERBB2*. We leveraged the TCGA cohort for discovery and the ICGC cohort along with a 341-patient cohort of ductal carcinoma *in situ* lesions (DCIS) for replication (**Figure 1D; Supplementary Table 1**). Briefly, we identified germline variants within ERBB2 that changed the protein sequence and predicted the number of epitopes derived from the variant protein sequence compared to WT sequence – defined as the reference genome. We confirmed genotype and HLA calls by ensuring allele frequencies in our cohort matched those reported in Gnomad (**Supplementary Figure 1B**) and The Allele Frequency Net Database (**Supplementary Figure 1C**), respectively (*4, 36*). Patients were stratified by their GEB (*i.e.* greater than, the same as or less than WT) (**Supplementary Figure 1D**) and whether they were HER2+ or HER2-, as defined by PAM50 transcriptional signature (**Figure 1D**). If a high GEB in HER2 leads to increased immune surveillance upon *ERBB2* amplification, we hypothesized we would see a depletion of germline epitopes in HER2+ breast tumors compared to HER2- tumors. We discovered the GEB in *ERBB2* is significantly negatively associated with HER2+ breast cancer (OR = 0.59; P = 6.63x10^-3^; **Figure 1E**). The odds of developing HER2+ breast cancer was 38% lower if an individual has a high GEB in *ERBB2*. A similar association was observed when considering GEB as a continuous value (**Supplementary Figure 1E**). The association was robust to varying definitions of HER2 positivity (*i.e.* transcriptomic *vs* genomic amplification; **Supplementary Figure 1F**) and HLA binding (*i.e.* varying binding thresholds; **Supplementary Figure 1G**).

To ensure this association was not driven by functional germline variants that may promote or protect against HER2+ breast cancer, we reran our analyses holding germline variants constant but randomly reassigning the HLA alleles. If the observed associations are driven by germline variants alone, we should still see the same associations even with the null HLA alleles. However, if the presentation of the germline variants drives the observed depletion, the association should be abrogated with the null HLA alleles. Indeed, the negative association between GEB and HER2+ breast cancer was abrogated when HLA alleles were randomly assigned indicating that the association is not driven by functional germline variants alone (permutation test n = 1,000; **Supplementary Figure 1H**).

Next, we investigated if GEB selected against amplifications of other recurrently amplified genes in breast cancer (**Figure 1F**). The Integrative Cluster (IntClust) subtypes of breast cancer are characterized by distinct copy number and gene expression profiles (*32*). We focused on the four ER+/HER2- high-risk IntClust subtypes, IC1, IC2, IC6 and IC6, which are each defined by specific recurrent amplifications and have an elevated and persistent risk of recurrence up to two decades after diagnosis (*33*). Since TNBC is characterized by increased genomic instability rather than focal amplification of any one specific gene/region, it did not lend itself well to these analyses (*32*). For each of the four subtypes, we identified 3-4 recurrently amplified genes that were highly expressed (**Supplementary Figure 1I-L**) and calculated the GEB as the sum of the burden across the selected genes. To increase power, subtypes were defined by amplification of the defining cytoband and ER positivity (**Supplementary Figure 1M**). Similar to HER2, we found consistent negative associations between GEB and subtype membership (OR = 0.51-0.83; FDR = 0.033-0.68; **Figure 1G; Supplementary Table 1**). For example, a high GEB in *MYC, SQLE* and *FBXO32* was negatively associated with likelihood of developing the IC9 subtype which is defined by somatic amplification of these genes. Again, associations were depleted when HLA alleles were scrambled, indicating that these associations are not driven by functional variants alone (**Supplementary Figure 1H**). None of the variants considered in these analyses are known breast cancer susceptibility loci (**Supplementary Figure 1N**) (*7*) and GEB was not significantly correlated with somatic epitope burden in any of the subgroups (P > 0.48).

As negative controls, we identified proteins that were not expressed in mammary tissue as unexpressed proteins should not produce epitopes and, therefore, no association with subtype. We focused on keratins given their tissue specific expression patterns. We identified keratins that are not expressed in mammary tissue: KRT34, KRT71, KRT74 and KRT82 (*37, 38*) and tested the association between the GEB with the four PAM50 subtypes (excluding normal-like; **Supplementary Figure 1O**). None of the keratins were significantly associated with any subtype and the effects of the subtype-specific genes were significantly stronger than the keratin negative controls (**Supplementary Figure 1P**). Taken together, these data indicate that germline-mediated immunoediting sculpts the molecular subtype a tumor commits to.

### Germline-mediated immunoediting dictates breast cancer subtype during tumorigenesis

To investigate whether patterns of germline-mediated immunoediting generalize to other breast cancer cohorts and to elucidate the timing of this process during disease progression, we evaluated the association between GEB and subtype commitment in two additional cohorts: the ICGC cohort (n=431; **Supplementary Figure 2A**) (*31*) as well as a cohort of 341 ductal carcinoma *in situ* lesions (DCIS) subjected to shallow whole genome sequencing (sWGS) (*30*). Genotyping and ancestry estimation were conducted as described previously (*39*). Genotypes were confirmed for both cohorts by ensuring polymorphism and HLA allele frequency matched those reported in Gnomad (**Supplementary Figure 2B-C**) and The Allele Frequency Net Database (**Supplementary Figure 2D-E**), respectively (*4, 36*). Across all cohorts we observed consistent negative associations between GEB and subtype membership (**Figure 2A; Supplementary Figure 2F; Supplementary Table 1**). Meta-analysis indicates that the odds of developing any of the subtypes is 1.25-1.9 times lower if the individual harbors a high GEB (FDR = 0.047-0.12). Thus, these data demonstrate that high GEB selects against subtype commitment and this negative selection is observed early, during pre-neoplasia.

**Figure 2.**
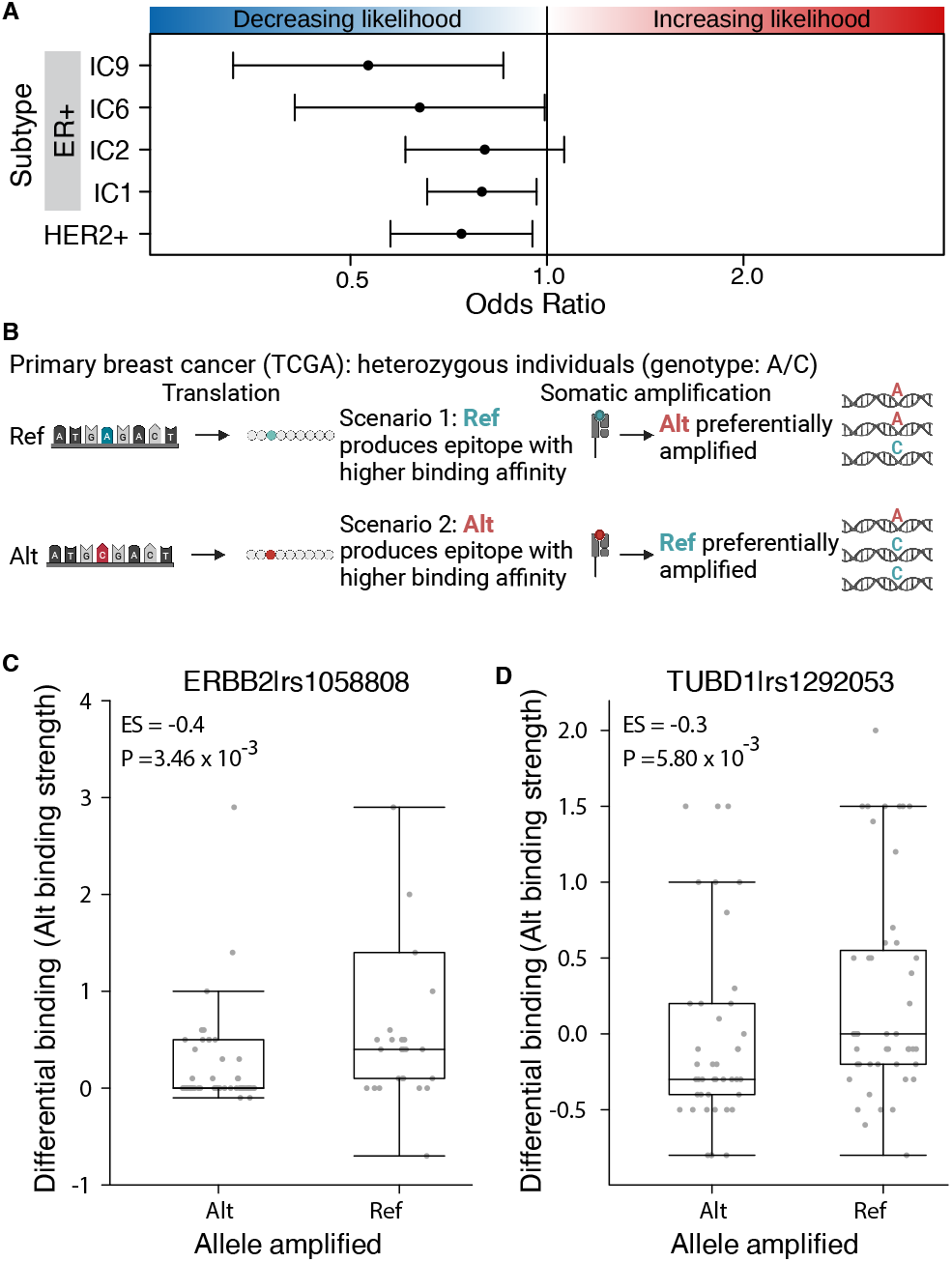
Germline-mediated immunoediting dictates breast cancer subtype early during tumorigenesis. **A)** Across five subtypes and three independent cohorts, high GEB in subtype-specific oncogenes is associated with a decreased likelihood of developing the cognate subtype of breast cancer. Forest plot shows the odds ratio and 95% confidence intervals from a meta-analysis across three cohorts: DCIS (N=341), TCGA (n=656) and ICGC (n=431). **B)** At the individual variant level, to avoid immunoediting, tumors should preferentially amplify the germline allele that produces a weaker epitope. For example, considering only heterozygous individuals, in scenario 1 the reference (ref) allele produces an epitope with higher binding affinity for MHC class I than the alternative (alt) allele. As a result, the tumor preferentially amplifies the alt allele as evidenced by the increased proportion of sequencing reads supporting the alt allele. By contrast, in scenario 2, the alt allele produces an epitope with higher binding affinity for MHC class I and the ref allele is preferentially amplified. **C-D)** Allele producing epitope with weaker MHC class I binding affinity is preferentially amplified. Boxplots of differential binding for epitopes derived from the alt allele *vs* ref allele (y-axis), *i.e.* a measure of alt allele binding affinity, for samples that preferentially amplified the alt or the ref allele. Effect size and p-value from Mann-Whitney rank sum test. Boxplots show analysis for rs1058808 derived from *ERBB2* **(C)** and rs1292053 derived from *TUBD1* **(D)**.

Next, we investigated if germline-mediated negative selection persists in metastatic lesions. To this end, we leveraged a fourth cohort of 702 metastatic breast cancer tumors with blood and tumor whole genome sequencing (*29*) (**Figure 1D**). Once again, we confirmed genotypes according to Gnomad and The Allele Frequency Net Database (**Supplementary Figure 2G-H**) (*4, 36*) and considered individuals of European descent (**Supplementary Figure 2I**). The association between GEB and subtype commitment was significantly weaker in metastatic *vs.* primary tumors and DCIS or, in some cases, the directionality changed (**Supplementary Figure 2J; Supplementary Table 1**). These data suggest immune sculpting occurs early during breast tumorigenesis and by the time the tumor metastasizes, it may no longer be susceptible to immunoediting pressures (*40*).

### MHC class I binding affinity influences allele specific amplification

If germline epitopes are involved in immunoediting, we should observe preferential gene amplification at loci harboring heterozygous polymorphisms. Suppose one allele produces an epitope with a higher MHC class I binding affinity than the alternative allele. In that case, the allele with the lower binding affinity will be preferentially amplified as it elicits a weaker immune response and has a lower fitness cost associated with its amplification (**Figure 2B**). To test this, we focused on the two most common variants observed in our discovery cohort: rs1058808 in *ERBB2* (MAF = 0.33) and rs1292053 in *TUBD1* (MAF = 0.45). For both variants, we identified individuals that were heterozygous with either *ERBB2* or *TUBD1* amplifications, respectively. Next, we identified individuals that preferentially amplified the reference (Proportion ReadsAlt < 0.20) or the alternative allele (Proportion ReadsAlt > 0.80; **Supplementary Figure 2K-L**). For each individual, we calculated the binding affinity of the alternative allele, defined as the differential binding affinity of the alternative allele compared to the reference allele (Altbinding – Refbinding; **Figure 2B**). A high differential binding indicates the epitope derived from the alternative allele outcompetes the reference allele. For both rs1058808 and rs1292053, we observed significantly higher differential binding – *i.e.* alt allele binding affinity – in individuals that preferentially amplified the reference allele over the alternative allele (ESrs1058808 = -0.4; Prs1058808 = 3.46x10^-3^; ESrs1292053 = -0.3; Prs1292053 = 5.800x10^-3^; Mann-Whitney rank sum test; **Figure 2C-D**). We confirmed these data were not driven by any one tumor by testing the median differential binding per sample (ESrs1058808 = -0.4; Prs1058808 = 1.34x10^-2^; ESrs1292053 = -0.25; Prs1292053 = 1.59x10^-2^; Mann-Whitney rank sum test; **Supplementary Figure 2M-N**). These data suggest that tumors preferentially amplify the allele producing the epitope with weaker MHC class I binding affinity, consistent with our hypothesis that germline epitopes select against amplification.

### Tumors that overcome high GEB are more aggressive

As stated above, our data indicate that by the time the tumor metastasizes it may no longer be susceptible to immunoediting (**Supplementary Figure 2J**). The lack of evident germline-mediated immune editing in metastatic lesions suggests that at least a subset of tumors acquire oncogene amplification despite high GEB. Within each subtype, we investigated whether GEB was different between individuals with primary breast cancer *vs.* those with metastatic disease (**Figure 3A**). We first ensured there were no differences in allele frequencies for either the germline variants or the HLA alleles between the metastatic and primary tumors (**Supplementary Figure 3A-D**). Next, we compared the *ERBB2* GEB in HER2+ primary breast tumors to HER2+ metastatic tumors. The GEB was significantly higher in HER2+ metastatic tumors *vs.* primary tumors from two independent cohorts (TCGA: OR = 1.66; P = 0.076; ICGC: OR = 3.14; P = 0.033; **Figure 3B; Supplementary Figure 3E; table S1**). This observation extended to the ER+ high risk IntClust subtypes. For all four high risk IntClust subtypes, metastatic tumors were enriched for germline-derived epitopes in subtype-specific genes compared to primary tumors from both TCGA and ICGC (OR = 1.29-2.53; FDR < 0.22; meta-analysis; **Figure 3B; Supplementary Figure 3E**). No enrichment was observed when running the same analyses with scrambled HLA alleles indicating that the observed enrichment is not driven by germline variants alone but rather MHC class I presentation of germline-derived epitopes (**Supplementary Figure 3F**). These data suggest that within the same subtype, tumors that amplify a specific oncogene, despite a high GEB, have a propensity to be more aggressive than tumors with a low GEB in the amplified oncogene.

**Figure 3.**
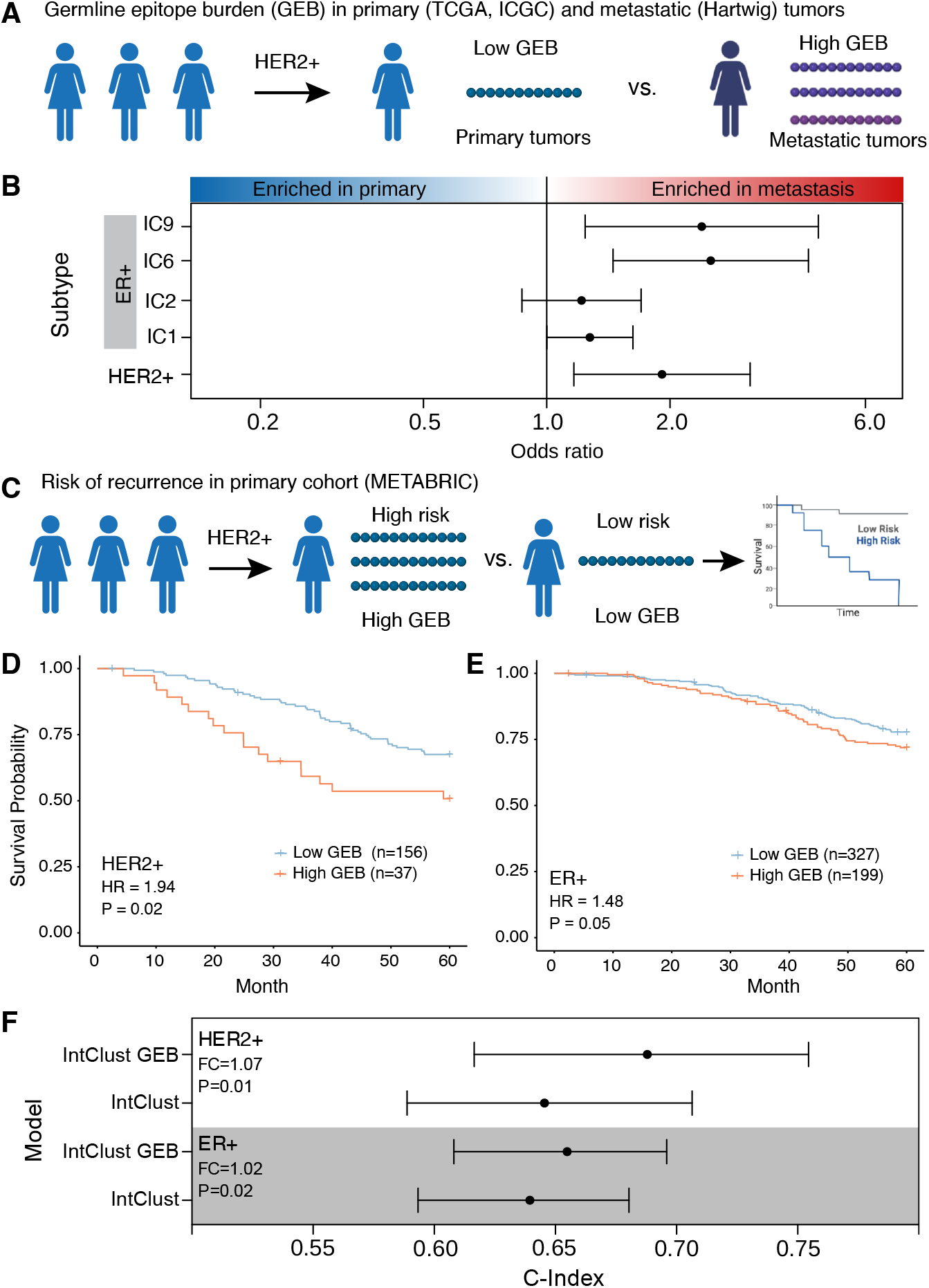
Tumors that overcome a high burden of germline epitopes are more aggressive. **A)** Schematic of within-subtype comparison between GEB in primary tumors (TCGA and ICGC) *vs* metastatic tumors (Hartwig). **B)** Across five subtypes, metastatic tumors show an enrichment of epitopes compared to primary tumors. Forest plot shows odds ratio and 95% confidence intervals from meta-analysis of TCGA *vs* Hartwig and ICGC *vs* Hartwig. **C)** Schematic of within-subtype comparisons of GEB association with risk of relapse within five years in METABRIC. **D-E)** A high GEB is associated with increased risk of relapse in HER2+ **(D)** and ER+ **(E)** tumors. Hazard ratio and p-value from CoxPH model correcting for first two genetic principal components, percent genome altered and age. **F)** GEB in combination with the Integrative Clusters (IntClust) improves the accuracy of five-year relapse prediction in ER+ and HER2+ tumors. Forest plot shows c-index of predictive models considering the IntClust alone or in combination with GEB for 1,000 bootstrapped iterations. Fold change (FC) is calculated as the ratio of medians while the p-value is calculated as 1 – the proportion of iterations where the -index of the IntClust and GEB model was greater than the IntClust alone model.

To strengthen the link between GEB and disease aggression, we investigated the association with disease recurrence (distant relapse) after adjuvant treatment of primary tumors (**Figure 3C**). To ensure sufficient power, we leveraged a fifth cohort, METABRIC, comprised of 1,725 primary breast cancer tumors. While METABRIC provides the largest primary breast cancer cohort with comprehensive 20 year follow-up, it was profiled with Affymetrix SNP 6.0 arrays which makes genotyping rare variants and HLA alleles challenging. Despite this limitation, benchmarking HLA imputation from SNP 6.0 arrays in TCGA, which has both SNP 6.0 and whole exome sequencing, identified a subset of HLAs with accurate (>80%) imputation (**Supplementary Figure 3G; see Methods**). Focusing on the subset of HLA alleles with accurate imputation, we tested the association between GEB and risk of relapse within the first 5-years as we hypothesize immunoediting happens early during tumor growth. We found that high GEB was associated with an increased risk of relapse within 5-years in HER2+ tumors (HR = 1.94; P = 0.02; CoxPH model; **Figure 3D; Supplementary Table 1**). To improve power, we considered the four ER+ high risk subtypes together, *i.e.* IC1, IC2, IC6 and IC9, and scored the GEB across all genes weighted by the presence of an amplification (see **Methods**). The high risk ER+ tumors showed the same trend – high GEB is associated with an increased risk of relapse within 5-years (HR=1.48; P = 0.05; **Figure 3E; Supplementary Table 1**). The same prognostic trends were observed within each subtype individually (**Supplementary Figure 3H**). The breadth of neoantigens that an HLA can present (*i.e.* HLA promiscuity) has previously been shown to be negatively associated with outcome following checkpoint blockade (*41*). To determine if the prognostic associations observed here were driven by HLA promiscuity, we calculated the proportion of promiscuous HLAs per individual (see **Methods**). While the proportion of promiscuous HLA alleles was weakly associated with outcome in HER2+ and high ER+ tumors (HRHER2= 3.99; PHER2= 0.071; HRER=2.15; PER=0.21; CoxPH model), controlling for HLA promiscuity did not abrogate the prognostic value of GEB (HRHER2= 1.81; PHER2= 0.036; HRER=1.50; PER=0.04; CoxPH model), suggesting this does not drive the observed prognostic associations.

Previously, we demonstrated that the genomically defined IntClust subgroups dramatically improve risk of relapse prediction in ER+/Her2- disease beyond the established clinical covariates, and this is especially true for late (>5 years) distant relapse (*33*). Next, we investigated whether GEB might improve relapse prediction. GEB in combination with the IntClust subgroups showed significantly higher relapse prediction accuracy than the IntClust subgroups alone (FC > 1.03; P < 0.03; **Figure 3F**). A more modest increase in accuracy was observed when considering clinicopathologic features in the model, including age, node involvement, size and grade (**Supplementary Figure 3I**). Taken together, these data suggest that tumors that overcome high GEB are aggressive. Moreover, GEB may improve predictions of risk of relapse in ER+ and HER2+ breast cancer.

### High GEB associated with lymphocyte depletion

In invasive breast cancer, the association between high GEB and a more aggressive tumor is counterintuitive as an increased number of epitopes should make the nascent tumor more detectable by the immune system. We reasoned that tumors with high GEB are forced to develop immune suppression or evasion mechanisms in order to survive. To test this hypothesis, we harnessed a previous immunogenomic characterization of TCGA breast tumors in which comprehensive immune-related transcriptional signatures and deconvolution methods were deployed on bulk RNA-sequencing to characterize the tumor microenvironment (*42*). Leveraging this resource, we identified immune cell population estimates, transcriptional signatures and gene markers reflective of immune cell infiltration, cytokine signaling and extracellular matrix composition (**Figure 4A; Supplementary Table 2**). Unsupervised clustering of these immune features in HER2+ tumors identified two clusters (**Figure 4B**). Cluster 1 was characterized by macrophages and TGFβ signaling, while cluster 2 was characterized by lymphocytes and interferon gamma signaling. Supportive of our hypothesis, the myeloid predominant cluster 1 was enriched for HER2+ tumors with high GEB while the lymphocyte predominant cluster 2 was enriched for HER2+ tumors with a low GEB (OR = 4.64; P = 0.012; **Figure 4B**). This enrichment was confirmed with consensus clustering (OR=3.25; P = 0.051; **Figure 4B**). Similarly, clustering of ER+ tumors identified a lymphocyte predominant cluster enriched for tumors with a low GEB and a myeloid predominant cluster enriched for tumors with high GEB (OR = 2.56; P = 0.02; **Figure 4C**). Next, we interrogated lymphocyte and macrophage infiltration in high *vs.* low GEB tumors. In both HER2+ and ER+ tumors, we observed a decrease in lymphocytes in tumors with high GEB (Effect Size ≤ -0.05; FDR ≤ 0.10; **Supplementary Figure 4A-B**). A modest downregulation of cytotoxicity, defined as the geometric mean of *GZMA* and *PRF1* abundance, was also observed (Effect Size ≤ -21.71; FDR ≤ 0.16; Mann-Whitney test; **Supplementary Figure 4C**). In parallel, tumors with high GEB demonstrated higher macrophage infiltration, specifically M2- rather than M1-polarized macrophages (Effect Size ≥ 0.06; FDR ≤ 0.10; Mann-Whitney test; **Supplementary Figure 4D-F**). These data are consistent with high GEB tumors having increased macrophage and lower lymphocyte infiltration compared to low GEB tumors, regardless of subtype. Next, we considered the six immune subtypes identified by Thorsson *et al* (*42*). We asked if, within each breast cancer subtype, the GEB was associated with anti-tumor (*e.g.* IFN-*γ* dominant, inflammatory) or tumor-promoting (*e.g.* wound healing, lymphocyte depleted, immunologically quiet, TGF-β dominant) immune microenvironments. In both HER2+ and ER+ tumors, high GEB was associated with tumor-promoting immune subtypes, although the associations did not reach statistical significance (ORHER2+ = 0.50; PHER2+ = 0.14; ORER+ = 0.70; PER+ = 0.24; **Supplementary Figure 4G**). Taken together, these data demonstrate that tumors which overcome a high GEB have less cytotoxic immune microenvironments.

**Figure 4.**
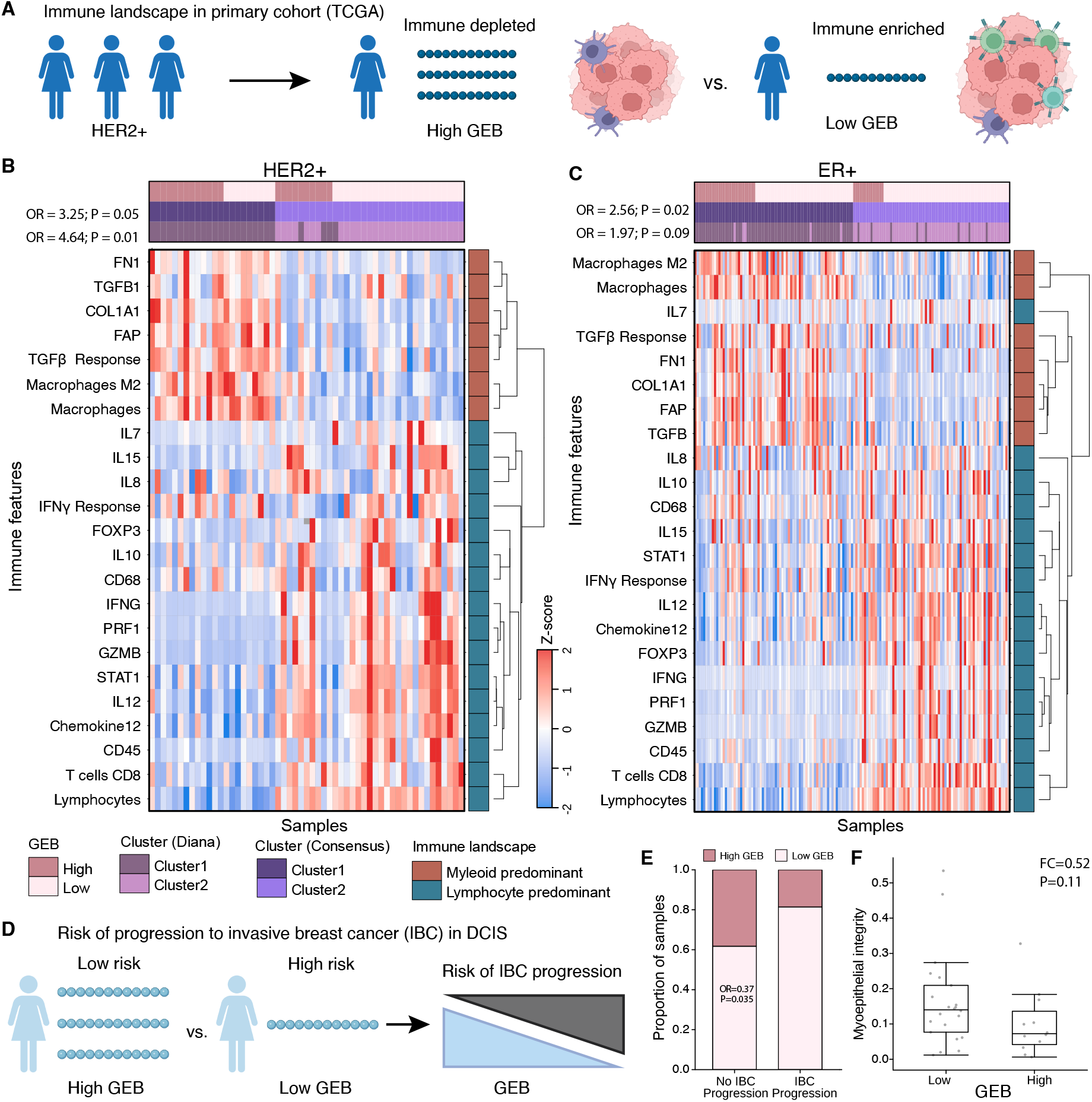
A high germline epitope burden promotes an immunosuppressive phenotype. **A)** Schematic of within-subtype comparisons of the immune landscape between high GEB and low GEB tumors in TCGA. **B-C)** Unsupervised clustering of 23 immune features, selected to reflect broad immune cell populations, cytokine signaling and extracellular matrix composition, identified two dominant clusters within HER2+ **(B)** and ER+ **(C)** breast tumors driven by GEB. Heatmap shows the z-score of each immune feature (y-axis) for each tumor (x-axis). Covariates along the top indicate if the tumor has a high GEB or a low GEB along with clusters from two different clustering methods (Diana and Consensus). Immune features cluster into two broad categories, myeloid and lymphocyte predominant, as indicated by the covariate on the right. Statistics from Fisher’s exact test quantifying the enrichment of high GEB tumors in the myeloid predominant cluster for both clustering methods. **D)** Schematic of GEB association with progression to invasive breast cancer. **E)** DCIS lesions that do not progress to IBC are enriched for high GEB. Barplot shows the proportion of DCIS lesions that progress or not progress to IBC stratified by GEB. Statistics from a logistic regression model correcting for eight genetic principal components, HER2 and ER status. **F)** Myoepithelial integrity is negatively associated with GEB. Boxplot shows myoepithelial integrity (% of E-cadherin in myoepithelium), as defined by Risom et al. (*43*), in high vs low GEB lesions for a subset of lesions that had spatial proteomics data. Statistics are based on a Mann-Whitney Rank Sum test.

Finally, we asked whether the observed differences in the tumor immune landscape could be explained instead by disruption to MHC class I presentation, a recurrent mechanism of immune evasion (*40*). We considered two transcriptional signatures of MHC class I presentation, as evaluated by Thorsson et al. (*42*). In both HER2+ and ER+ tumors, we observed only a slight decrease in MHC class I presentation in high GEB compared to low GEB tumors (ESER+ = -0.01; PER+ = 0.06; ESER+ = -0.02; PER+ = 0.07; Mann-Whitney test; **Supplementary Figure 4H-I**). This modest trend suggests that downregulation of MHC class I presentation may be one mechanism by which tumors with a high GEB evade immune detection but that it is unlikely to drive the observed immune differences.

### High GEB in DCIS reduces likelihood of progression to invasive breast cancer

The model of germline-mediated immunoediting posits that, prior to immune escape, tumors with a high GEB are more likely to undergo immune-mediated cell death. Immune selection pressures promote immune escape resulting in invasive breast cancers (IBC) with high GEB being more aggressive. We reasoned that DCIS likely represents an immune protected state with more susceptibility to immunosurveillance, *i.e.* a state prior to immune escape. We hypothesized that high GEB in DCIS would be associated with decreased likelihood of progression to IBC, opposite to that observed in primary IBC (**Figure 4D**). In line with our hypothesis, DCIS lesions that did not progress to IBC were enriched for high GEB (OR = 0.37; P = 0.035; **Figure 4E**). Previously, Risom et al. demonstrated that myoepithelial disruption was more pronounced in DCIS lesions that did not progress to IBC (*43*). Similarly, in a small cohort of the DCIS lesions with spatial proteomics data (n=34), we observe a modest negative association between GEB and myoepithelial integrity consistent with high GEB lesions experiencing more immune surveillance (FC = 0.52; P = 0.11; **Figure 4F**). Taken together, these data indicate DCIS lesions with a high GEB may experience more immune surveillance and GEB may be predictive of progression to invasive breast cancer. In order to progress to IBC, lesions must suppress or escape immune surveillance resulting in more aggressive primary tumors with a proclivity to metastasize.

Altogether, our findings demonstrate that the germline genome influences somatic evolution *via* immunoediting. If a nascent lesion develops in an individual with high GEB in key breast cancer associated oncogenes, somatic amplification of the cognate oncogene comes with a fitness cost to the tumor (**Figure 5**). Oncogene amplification further augments an already high GEB, thereby increasing the likelihood of immune surveillance. Thus, the lesion is less likely to acquire a somatic amplification in the oncogene of interest, instead favoring an alternative oncogenic pathway with lower fitness costs. If the lesion is able to overcome high GEB and still acquires somatic amplification of the oncogene, the tumor is more aggressive and has a lymphocyte-depleted tumor microenvironment. This new model of germline-mediated immunoediting can be exploited to refine risk stratifications within breast cancer subtypes. Within subtypes already defined as part of the treatment paradigm of breast cancer, individuals with a high GEB and, therefore, an increased risk of relapse can be identified from germline information measured from blood. In this way, GEB may represent a minimally invasive biomarker to further refine breast cancer relapse risk and accompanying treatment and monitoring.

**Figure 5.**
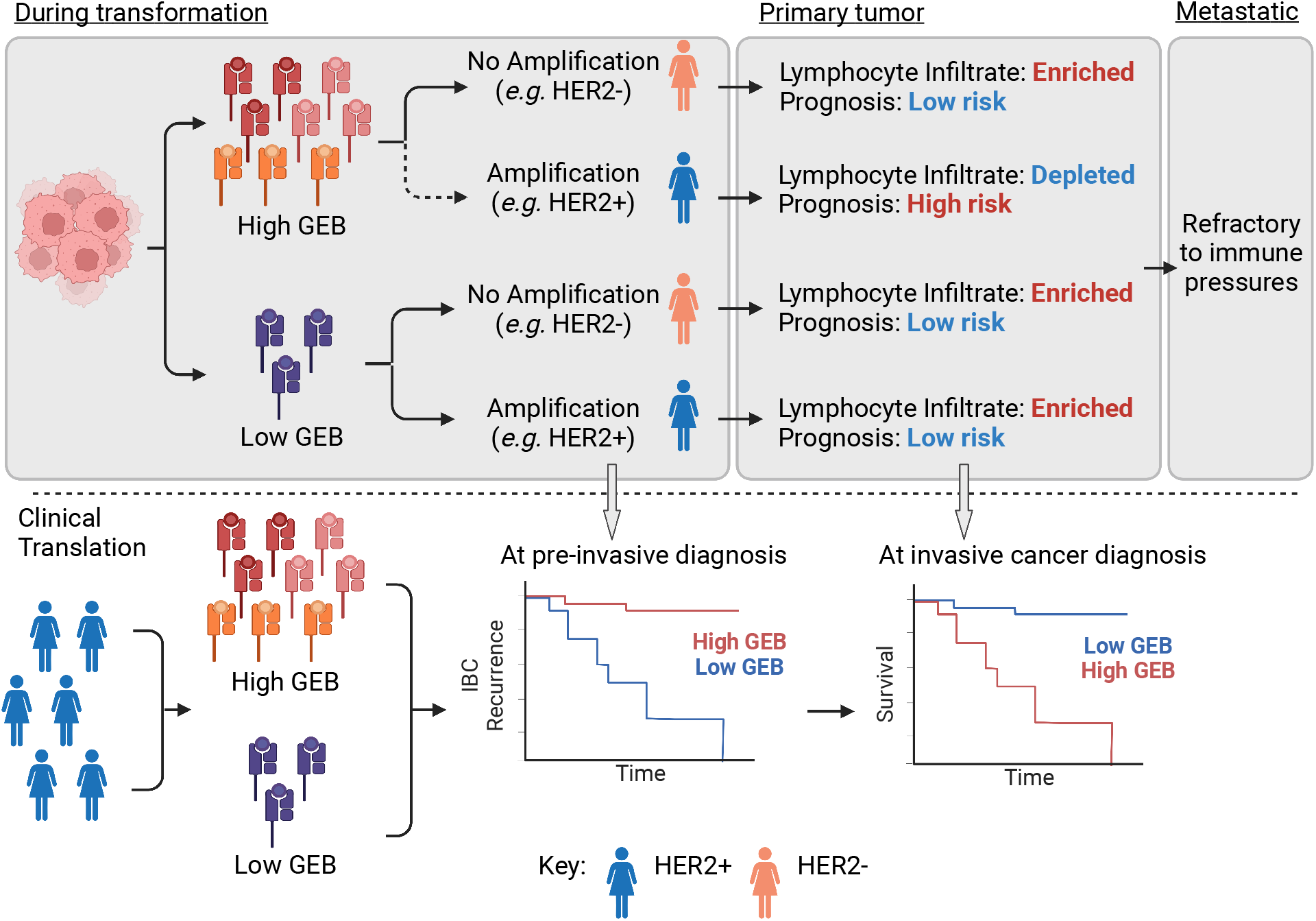
Germline-mediated immunoediting sculpts breast cancer subtypes and metastatic proclivity. Schematic of mechanistic molecular model of germline-mediated immunoediting and its implications for improving breast cancer risk stratification. Briefly, during tumorigenesis, lesions with a high GEB in a gene of interest are less likely to acquire somatic amplification of that gene. However, if the tumor gains additional copies of the gene, it is forced to develop an immune suppressive/evasive phenotype and is more aggressive. Conversely, low GEB has little impact. By the time the tumor has metastasized to distant sites, it develops immune suppression/evasion mechanisms and is refractory to immunoediting pressures. In the pre-cancerous setting, GEB may be indicative of risk of progression to an invasive cancer since lesions with high GEB would have to overcome stronger immune pressures. Once the lesion becomes invasive, within a breast cancer subtype, tumors may be further stratified into those with high and low risk of relapse based on GEB in subtype-specific genes.

## Discussion

Leveraging 3,855 breast cancer lesions, spanning pre-invasive to metastatic disease, we show that a high GEB in recurrent oncogenes is associated with a reduced likelihood of somatic amplification of the oncogene (**Figure 5**). We reason this is because somatic amplification of a gene with a high GEB would increase the number of available epitopes resulting in increased immunosurveillance. Some tumors are able to overcome this high GEB and amplify a particular oncogene; however, these tumors are more aggressive and have “immune cold”, lymphocyte depleted, microenvironments.

There is mounting evidence that germline variation can skew somatic mutagenesis. For example, loss-of-function variants in DNA repair genes, such as *BRCA1* and *BRCA2*, deregulate homologous repair resulting in mutational profiles characterized by tandem duplications and small (<10kb) deletions, respectively (*2*). Germline variants that regulate transcription of an oncogenic pathway can select for further somatic deregulation of the same pathway (*8*). Elucidating mechanistic links between germline variation and somatic mutagenesis facilitates delineation of the contribution of nature *vs.* nurture to inter-tumoral heterogeneity. We propose a novel mechanism by which the germline genome influences somatic evolution *via* immunoediting. Germline variants in a gene of interest that produce epitopes increases the fitness cost, thus decreasing the likelihood of gene amplification. Taken together, these studies suggest somatic mutagenesis is not stochastic but rather probabilistically bounded by the germline genome, which determines the likelihood of acquiring specific somatic mutations.

In addition to linking germline and somatic variation, these data elucidate a novel link between germline variation and the tumor immune landscape. Specifically, we show that breast tumors with high GEB in a particular oncogene that somatically amplify the oncogene are more likely to become “immune cold” tumors – decreased lymphocyte infiltration. With increasing evidence supporting the effectiveness of checkpoint blockade in the adjuvant setting in breast cancer, including FDA approval of Pembrolizumab as an adjuvant therapy in TNBC (*44*), refined biomarkers are needed to predict which tumors would benefit the most from this form of therapy. Moreover, various clinical trials are currently underway evaluating experimental vaccines in breast cancer. One such trial (NCT04367675) is investigating the safety profile of plasmid DNA vaccine encoding *hTERT*, *PMSA* and *WNT1* (INO-541) alone or in combination with IL12 (INO-9012) in BRCA1 or BRCA2 carriers (*45*). A second (NCT04674306) is investigating the safety and effective dose of an alpha-lactalbumin vaccine in triple negative breast cancer (*46*). Finally, GP2, a vaccine involving an endogenous peptide derived from HER2 (*34*), has progressed to a multi- center, randomized, phase three clinical trial to evaluate its safety and efficacy in HER+ breast cancer (NCT05232916). These studies further motivate the discovery of generalizable cancer vaccine targets to facilitate preventative vaccinations. Germline epitopes, such as those presented here, may serve as such generic targets.

Importantly, despite recent efforts to improve ancestry representation in cancer genomic cohorts, currently available cohorts are heavily enriched for individuals of European descent. Further studies in more diverse cohorts are required to evaluate the generalizability and prevalence of germline-mediated immunoediting in other populations, and may help to explain differences in breast cancer subtype and severity (*13*). While the current study focuses on breast cancer we anticipate germline-mediated immunoediting extends to other cancers. Future studies in adequately powered cohorts are needed to confirm this.

These data have important clinical implications. First, germline variants can be measured from blood and thus represent a low-cost, minimally invasive biomarker that is not sufficiently harnessed at present. Second, germline-mediated immunoediting may explain why an individual develops one breast cancer subtype over another. Third, GEB, measured from blood, may be exploited to further stratify risk of relapse within breast cancer subtypes as well as to identify tumors with high lymphocyte infiltration, *i.e.* “immune hot” tumors. Fourth, our data demonstrate that immunoediting pressures differ during the course of a patient’s disease, potentially informing the timing of therapeutic interventions. Finally, germline-mediated immunoediting points to a broad source of currently underappreciated immunogenic antigens, thus dramatically expanding the quest for alternative antigens (*47*). This motivates a potential new avenue for developing cancer vaccines with desired properties, including being clonal and present in entire subgroups of disease.

## Methods

### The Cancer Genome Atlas (TCGA) primary invasive breast cancer cohort

We considered 656 primary breast cancer tumors from TCGA that were determined to be of European descent by Yuan *et al* (*48*) and had both tumor and normal whole exome sequencing (*3*). Whole exome sequenced bam files were downloaded from Genomic Data Commons Data Portal (https://portal.gdc.cancer.gov/). We leveraged previously published copy number alterations, mRNA abundance and somatic single nucleotide variants downloaded from https://gdc.cancer.gov/about-data/publications/pancanatlas. Genes were considered to be amplified if the total copy number was greater than 4. We downloaded class 1 HLA alleles for each individual in TCGA from Shulka et al. (*49*).

### The International Cancer Genome Consortium (ICGC) primary invasive breast cancer cohort

We leveraged 431 primary breast cancer tumors with paired normal whole genome sequencing from ICGC (*31*). Bam files were downloaded from EGA (EGAD00001000141, EGAD00001001322, EGAD00001001334, EGAD00001001335, EGAD00001001336, EGAD00001001337, EGAD00001001338), backextracted using picard SamToFastq (v2.27.5), followed by alignment to hs37d5 using the PCAP (ICGC/TCGA Pan-Cancer Analysis Project) docker implementation of bwa mem (https://github.com/cancerit/PCAP-core). Genotypes were called using HaplotypeCaller (v4.1.8.1) to produce gvcfs, followed by joint genotyping (*i.e.* GenotypeGVCFs) and variant recalibration (*i.e.* VariantRecalibrator and ApplyVSQR), according to GATK best practices (*50*). Using 128 ancestry informative markers (*51*), we estimated the genetic ancestry of each case *via* principal component comparison with reference populations provided by Philips *et al*. Cases that clustered outside +/- 0.2 from the European-descent population in the reference population were excluded from further analyses. We leveraged previously published copy number segments provided in Supplementary Table 4 of Nik-Zainal et al. (*31*). Genes were considered to be amplified if the total copy number was greater than 4. Hormone status was provided in Supplementary Table 1 of Nik-Zainal *et al*. We identified MHC class I alleles using the nf-core/hlatyping nextflow (v21.05.0) pipeline (revision 1.2.0) using default parameters.

### GP2 presentation selects against HER2+ breast cancer

GP2 is a peptide vaccine derived from HER2 with known immunogenicity. To evaluate if GP2 presentation was associated with an individual’s likelihood of developing HER2+ breast cancer, we first identified HLA alleles that could present GP2 using netMHCpan (v4.1) with default settings (*52*). We tested all HLA alleles and considered those where GP2 was ranked within the top 3% of peptides – the same threshold as used throughout the manuscript (see **Germline-derived epitope burden in TCGA**). Next, we calculated the number of HLA alleles each individual possessed that were capable of binding GP2. We tested if the number of HLA alleles was associated with HER2+ breast cancer, defined by clinical hormone status, using a logistic regression model correcting for the first six genetic principal components:

HER2+ ∼ number of HLA alleles able to bind GP2 + PC1 + PC2 + PC3 + PC4 + PC5 + PC6

The same model was applied to both TCGA and ICGC. Meta-analysis was conducted using restricted maximum likelihood (REML) as implemented in the metafor (v3.4.0) R package.

### Germline-derived epitope burden in TCGA

We identified genomic coordinates representing recurrently amplified genes in the high-risk ER+/HER2- integrative subtypes (IntClust: IC1, IC2, IC6, IC9) and HER2+ (IC5) breast cancer subtypes (*32, 33*) (**Figure 1F**). We selected the three or four most abundant genes based on mRNA profiling for each high-risk ER+ amplicon using the RNA supplemental data provided as part of the TCGA PanCancer Atlas (*3*): IC1 – *RPS6KB1, TUBD1*, *DHX40, BCAS3;* IC2 – *RSF1, CCND1, PAK1, NARS2*; IC6 – *ZNF703, FGFR1, LETM2*, *EIF4EBP1*; IC9 – *MYC, SQLE, FBXO32*. Using these coordinates, we ran GATK HaplotypeCaller (v4.1.8.1) to produce gvcfs, followed by joint genotyping (*i.e.* GenotypeGVCFs) and variant recalibration (*i.e.* VariantRecalibrator and ApplyVSQR), according to GATK best practices (*50*). For each gene, we identified missense and frameshift variants using SnpEff (v4.3t) and generated the protein sequence considering all missense and frameshift variants using FRED-2 (v2.0.7) (*53*). Next, in order to calculate how many epitopes could be derived from each protein sequence, we downloaded class 1 HLA alleles for each individual in TCGA from Shulka *et al* (*49*). We predicted the binding probabilities for all 8- 11 amino acid peptides derived from each gene for each class I HLA allele using netMHCpan (v4.1) with default settings (*52*). For each individual, considering their class I HLA alleles, we also predicted the binding probabilities for peptides derived from the reference sequence for each gene using the same approach. To enrich for peptides able to escape central tolerance (*21*), we considered peptide sequences that were predicted to be weak binders. We empirically evaluated various binding thresholds before defining “weak binders” as within the 0.5-3% of naturally occurring random peptides (**Supplementary Figure 1G**). We calculated the total number of weak binding peptides derived from each individual’s protein sequence, including all missense and frameshift germline variants, and subtracted the total number of weak binding peptides derived from the reference protein sequence. This resulted in a value for each of the six class I alleles for all isoforms of the gene of interest. Next, for each class I allele, we took the median over all isoforms. Finally, we calculated the median over all six class I alleles. This resulted in the average GEB for each gene for each individual with respect to the reference sequence. We binned samples with GEB less than the reference sequence (< WT), the same as the reference sequence (WT) or more than the reference sequence (>WT). When considering multiple genes, we first binned samples with epitope burden <WT (-1), WT (0) or >WT (1) per gene first before summing across genes.

### Germline-derived epitope burden in ICGC

Genotypes were called using HaplotypeCaller (v4.1.8.1) to produce gvcfs, followed by joint genotyping (*i.e.* GenotypeGVCFs) and variant recalibration (*i.e.* VariantRecalibrator and ApplyVSQR), according to GATK best practices (*50*). We identified MHC class I alleles using the nf-core/hlatyping nextflow (v21.05.0) pipeline using default parameters. HLA allele frequencies within the ICGC cohort were compared against population frequencies from the Allele Frequency Net Database considering USA NMDP European Caucasian populations. Calculations of the average GEB were performed as for the discovery cohort (see **Germline-derived epitope burden in TCGA**).

### Ductal carcinoma *in situ* (DCIS) cohort

We leveraged 341 primary ductal carcinoma in situ tumors shallow whole genome (sWGS) sequencing from HTAN Atlas (*30*). Samples were sequenced and sequencing data aligned as previously described (*30*). Briefly, raw reads were aligned to GRCh38 reference genome using BWA (v0.7.17) and GATK (v4.1.7.0) (*54*) implemented in the Nextflow-base pipeline Sarek (v2.6.1). The recalibrated reads were further processed and filtered for mappability, GC content using the R/Bioconductor quantitative DNA-sequencing (QDNAseq) (v1.22.0) with R (v3.6.0). Copy number alterations were called using QDNAseq as detailed in Strand et al. (*30*). A subset of lesions (n=34) additionally had multiplexed ion beam imaging as previously described (*43*).

### Germline-derived epitope burden in DCIS

We used QUILT (v0.1.9) to simultaneously genotype and impute germline variants genome-wide (*55*). We used the 1000 Genomes Project as the reference panel and the following parameters “-- buffer=10000 --nGen=100”. QUILT was also used to estimate the class I HLA alleles for each case leveraging the reference panels provided with QUILT and default parameters. We only considered samples that had more than two HLA alleles imputed with >50% posterior probability (n=341 samples). Calculations of the average GEB were performed as for the discovery cohort (see **Germline-derived epitope burden in TCGA**).

### Germline-derived epitope burden in metastatic breast cancer (Hartwig)

We leveraged 702 metastatic breast cancer tumors with paired normal whole genome sequencing from Hartwig (*29*). Genotypes were called using HaplotypeCaller (v4.1.8.1) to produce gvcfs, followed by joint genotyping (*i.e.* GenotypeGVCFs) and variant recalibration (*i.e.* VariantRecalibrator and ApplyVSQR), according to GATK best practices (*50*). Using 128 ancestry informative markers (*51*), we estimated the genetic ancestry of each case *via* principal component comparison with reference populations provided by Philips *et al*. Cases that clustered outside +/- 0.2 from the European-descent population in the reference population were excluded from further analyses. We identified MHC class I alleles using the nf-core/haplotyping nextflow (v21.05.0) pipeline (revision 1.2.0) using default parameters. HLA allele frequencies within the Hartwig cohort were compared against population frequencies from the Allele Frequency Net Database considering USA NMDP European Caucasian populations. Calculations of the average GEB were performed as for the discovery cohort (see **Germline-derived epitope burden in TCGA**). Copy number alterations were identified using FACETS-SUITE with the following parameters: “--cval 1000 --purity-cval 1800 --normal-depth 20”. A gene was considered amplified is if the total copy number was greater than 4.

### Association between germline-derived epitope burden and oncogene amplification

We hypothesized that a high GEB in an oncogene would select against oncogene amplification. To test this, we considered five breast cancer subtypes defined by amplification of characteristic genomic regions: HER2+ and *ERBB2* (17q12), IC1 and 17q23, IC2 and 11q13, IC6 and 8p12, and IC9 and 8q24. To identify which genes within these amplified regions to measure germline epitope burden, we prioritized genes with high mRNA abundance. Specifically, we selected four genes with the highest median mRNA abundance per subtype based on RNA sequencing data provided by the TCGA PanCancer Atlas (**Supplementary Figure 1I-L**). Only three genes were selected for IC9 as only four genes are commonly amplified in this subgroup and *ADCY8* was not expressed (**Supplementary Figure 1L**). GEB across the genes selected was calculated as the sum of the sign of the epitope burden per gene, *i.e.* <WT (-1), WT (0) and >WT (1). We calculated the association between the GEB in each gene and whether each individual developed the corresponding subtype using a logistic regression model correcting for the first six genetic principal components and the somatic mutation burden:

subtype ∼ epitopes + PC1 + PC2 + PC3 + PC4 + PC5 + PC6 + somatic SNV burden

For TCGA, the first six genetic principal components calculated by Yuan et al. were used (*48*). For ICGC and Hartwig, genetic principal components were derived from PCA with reference populations provided by Philips *et al*. (*51*). Further modifications to the model were made for the DCIS cohort for which tumor tissue only was profiled with shallow whole genome sequencing (sWGS). Specifically, we only included HLA alleles genotyped with a posterior probability > 0.5. If after removing low confidence HLA alleles from the analysis, the sample had at most two unique HLA alleles remaining, we excluded the sample from the analysis. Additionally, we estimated ancestry in the sWGS DCIS cohort using the same protocol as Gusev et al. (*39*). Specifically, we scored each sample against ancestry-specific scores derived from SNPWEIGHTS (*56*) as implemented on github (https://github.com/gusevlab/panel-imp). We used these ancestry-specific scores to correct for population structure in the logistic regression model. As these samples were profiled with sWGS, we were not able to control for somatic SNV burden in the DCIS cohort. Meta-analysis was conducted using REML as implemented in the metafor (v3.4.0) R package. Finally, to improve power and unify subtype definitions across cohorts, subtypes in each cohort were defined as follows: HER2+ was defined as overexpression of HER2/*ERBB2* by PAM50 (TCGA, DCIS) or hormone status included in clinical annotations (ICGC, Hartwig), IC1 was defined as amplification of *RPS6KB1* and positive ER hormone status included in clinical annotations (ER+), IC2 was defined as amplification of *RSF1* and ER+, IC6 was defined as amplification of *ZNF703* and ER+ and IC9 was defined as amplification of MYC and ER+. P-values were adjusted for multiple hypothesis testing using the Benjamini-Hochberg correction. We also correlated GEB with somatic neoantigens as calculated by Thorsson et al. using Spearman’s correlation (*42*) .

### Null association between germline-derived epitope burden and oncogene amplification

To ensure the negative association observed between the GEB and oncogene amplification was not driven by germline variants having a functional impact on the gene itself, we permuted the HLA alleles, scrambling them across all individuals in the TCGA discovery cohort. We predicted epitopes using netMHCpan as described above using the scrambled HLA alleles instead of the true HLA genotypes. We calculated the average GEB for each gene and tested the association with the corresponding subtype using the null epitope predictions and the same modeling approach. We reran this permutation step 1,000 times and plotted the median, 0.025 and 0.975 quantiles of the null ϕ3 values.

### Allelic imbalance in somatic amplifications

To ensure sufficient power, we considered two of the most common SNPs in our discovery (TCGA) cohort: rs1058808 in *ERBB2* (MAF = 0.33) and rs1292053 in *TUBD1* (MAF = 0.45). For each variant, we identified heterozygous individuals that also had an amplification in the corresponding gene, *i.e. ERBB2* (defined as IC5) or *TUBD1* (defined as IC1). To determine allelic imbalance, we ran GATK CollectAllelicCounts (v 4.1.8.1) to calculate the number of reference and alternative allele at each site. We defined amplifications that preferentially amplified the reference allele as those that had <20% of reads mapping to the alternative allele. While amplifications that preferentially amplified the alternative allele were defined as those that had

>80% of reads mapping to the alternative allele. We ran the R package antigen garnish (v2.3.1) to identify the binding potential of peptides produced by the alternative allele (*57*). “Binders” were identified using the same definition as before: ranks within 0.5-3% of naturally occurring random peptides. We defined differential binding as the binding potential (*i.e.* rank) of the alternative allele – the reference allele. We compared differential binding between samples that preferentially amplified the alt allele *vs.* those that preferentially amplified the ref allele using a Mann-Whitney rank sum test. To ensure these differences were not driven by a single sample, we also evaluated the median differential binding per sample with preference to amplify the alt or ref allele.

### Germline epitope burden in primary vs metastatic tumors

We evaluated the GEB between primary and metastatic tumors within each subtype by applying a logistic regression model to compare the TCGA vs Hartwig cohorts and ICGC vs Hartwig cohorts separately:

metastatic (yes/no) ∼ epitopes

The model was applied to each subtype individually defining subtype as indicated previously (see **Association between germline-derived epitope burden and oncogene amplification)**. To ensure differences in GEB were not driven by differences in germline genotypes or HLA genotypes, we compared variant minor allele frequencies and HLA genotype frequencies using Pearson correlation. Meta-analysis was conducted using REML as implemented in the metafor (v3.4.0) R package. P-values were adjusted for multiple hypothesis testing using the Benjamini-Hochberg correction.

### Epitope burden across ER+ high risk tumors for within subtype analyses

To improve power for within subtype analyses, we merged all ER+ high risk tumors (*i.e.* IC1, IC2, IC6 and IC9). To ensure a uniform score was calculated across subtypes, we scored the GEB across all genes weighted by the binary presence of an amplification (defined as >4 copies):

ER+ high-risk GEB = IC1 genes burden * IC1 amplification + IC2 genes burden * IC2 amplification + IC6 gene burden * IC6 amplification + IC9 gene burden * IC9 amplification

This ensured the GEB within each set of subtype specific genes was only considered if the sample had the corresponding gene amplification.

### Benchmarking HLA imputation from SNP6.0 arrays

To assess the accuracy of HLA imputation from SNP6.0 arrays, we leveraged TCGA which has both SNP6.0 array and WES. We considered HLA alleles determined from WES previously as the gold standard (*49*). We called genotypes from SNP6.0 using Birdseed using default parameters and converted to plink file formats using plink (v1.9) (*58*). Next, we used CookHLA to impute HLA genotypes for each sample using the 1000 Genomes European reference panel (*59*). For each HLA allele, we assessed the concordance between the SNP6.0 imputed genotypes and the WES-derived gold standard genotypes. We only considered HLA alleles with >80% concordance (nalleles=68) for further analysis.

### Germline-derived epitope burden in METABRIC

We leveraged 1,725 primary breast cancer tumors with SNP6 arrays from METABRIC (*32*). Genotypes were imputed using the TOPMed Imputation Server using the TOPMed r2 reference panel and Eagle v2.4 for phasing (*60*). We used CookHLA to impute HLA genotypes for each sample using the 1000 Genomes European reference panel (*59*). Considering only HLA alleles with >80% concordance in our TCGA benchmarking (see **Benchmarking HLA imputation from SNP6.0 arrays**), we calculated the average GEB the same as the discovery cohort (see **Germline-derived epitope burden in TCGA**). We merged all ER+ high risk tumors (*i.e.* IC1, IC2, IC6, IC9) using the same scoring metric outlined in **Epitope burden across ER+ high risk tumors for within subtype analyses.**

### Prognostic association of germline-derived epitopes

We interrogated if GEB was associated with relapse after primary treatment in the METABRIC cohort (*32, 33*). Because we hypothesize immunoediting happens early in tumorigenesis, we focused on patients that relapsed within five-years of treatment. We tested prognostic associations in HER2+ and high risk ER+ tumors separately using a CoxPH model correcting for the first two genetic principal components, age at diagnosis, IntClust and percent genome copy-number altered (PGA). Within each subtype, samples were median dichotomized into low and high GEB. To test if the prognostic associations observed were driven by the capacity of each HLA to present a wide range of antigens (i.e. HLA promiscuity), we leveraged HLA promiscuity metrics generated by Manczinger et al. (*41*). For each sample, we calculated the proportion of HLA alleles that had a promiscuity score greater than the median promiscuity score. We only considered HLA alleles with scores measured by Manczinger et al. We tested the association between proportion of promiscuous HLAs and relapse using a CoxPH model correcting for the first two genetic principal components, age at diagnosis, PGA and IntClust subtype. Next, we tested if the proportion of promiscuous HLAs mediated the prognostic value of GEB using a logistic regression model including GEB, first two genetic principal components, age at diagnosis, IntClust, PGA and proportion of promiscuous HLAs. Finally, we evaluated if GEB could improve predictions of risk of five-year relapse. We first tested if GEB in combination with the IntClust subtypes improved relapse predictions over the IntClust subtypes alone. We compared the c-index of coxph models with GEB + IntClust subtypes *vs.* IntClust subtypes alone for 1,000 bootstrapped iterations. We calculated the fold change as the ratio of medians c-index from the two models and the p-value as 1 – the proportion of iterations where the c-index of the GEB + IntClust model was greater than the IntClust model alone. Next, we used the same approach to compare an IntClust and clinicopathologic model (IC + age + size + node involvement + grade) with IntClust, clinicopathologic and GEB model.

### Immune association of germline-derived epitopes

Within HER2+ and ER+ tumors, we interrogated the immune landscape of tumors with a high GEB *vs* tumors with a low GEB. We defined HER2+ based on pam50 and ER+ high risk tumors as IC1, IC2, IC6 and IC9 from the IntClust subtypes. We leveraged tumor microenvironmental characterization of the TCGA cohort conducted by Thorsson et al. (*42*). We selected 23 immune features – including cell estimates from deconvolution of RNA sequencing, transcriptomic signatures and marker gene mRNA abundance – based on their characterization of lymphocyte infiltration, macrophage infiltration, cytokine signaling and extracellular matrix composition. Immune features were downloaded from https://gdc.cancer.gov/about-data/publications/panimmune. We conducted unsupervised clustering based on Z-scores of the 23 immune features using Pearson correlation and the diana (DIvise ANAlysis Clustering) algorithm, as implemented in the cluster R package (v2.1.0), as well as consensus clustering as implemented in the ConsensusClusterPlus R package (v1.50.0). We compared how immune clusters associated with GEB using a Fisher’s exact test. We tested the association of lymphocyte and macrophage infiltration with GEB in HER2+ and ER+ tumors using a Mann-Whitney test.

Next, we considered the immune subtypes. We categorized subtypes into anti-tumor -- C2: IFN*γ* dominant and C3: inflammatory -- and pro-tumor -- C1: wound healing, C4: lymphocyte depleted, C5: immunologically quiet, C6: TGF-β dominant. We tested the association between the GEB and immune reactivity correcting for the first six genetic principal components:

Immune subtype ∼ GEB + PC1 + PC2 + PC3 + PC4 +PC5 +PC6

Finally, we investigated if differences in immune features were driven by disruption to HLA class I presentation. We considered two MHC class I antigen presentation signatures (Senbabaoglu - APM1 and Wolf - MHC.I_19272155) scored by Thorsson et al. (*42*) and downloaded from https://gdc.cancer.gov/about-data/publications/panimmune. Associations with GEB in HER2+ and ER+ tumors were quantified with a Mann-Whitney rank sum test. P-values were adjusted for multiple hypothesis testing using the Benjamini-Hochberg correction.

### Germline epitope burden association with risk of invasive breast cancer

To investigate if germline epitope burden was associated with risk of invasive breast cancer recurrence after DCIS, we calculated the germline epitope burden as the sum across 8p24, 8p12, 11q13, 17q23 and 17q12. High epitope burden tumors were defined as those with epitope burdens greater than the median. We evaluated if high germline epitope burden tumors were enriched in lesions that progressed to invasive breast cancer vs those that had neither invasive breast cancer nor DCIS recurrence using a logistic regression model correcting for eight genetic ancestry principal components, ER and HER2. Next, we identified 34 lesions that were also profiled with multiplex ion beam imaging (MIBI) (cite). Previously, myoepithelial integrity was negatively associated with risk of invasive breast cancer relapse. We evaluated if epitope burden was associated with myoepithelial integrity (defined as % E-cadherin within myoepithelial) using Mann-Whitney Rank Sum test.

### Data visualizations

Visualizations were generated in the R statistical environment (v3.3.1) with the lattice (v0.24-30), latticeExtra (v0.6-28) and BPG (v5.6.23) packages (*61*).

### Data availability

All cohorts are publicly available. TCGA BRCA samples can be found on the Genomic Data Commons Data Portal (https://portal.gdc.cancer.gov/). The ICGC breast cancer samples can be found on the European Genome-Phenome Archive (accession: EGAD00001000141, EGAD00001001322, EGAD00001001334, EGAD00001001335, EGAD00001001336, EGAD00001001337, EGAD00001001338). Metastatic breast cancer samples (Hartwig) are available for academic use under a Data Use Agreement (DR-230) from the Hartwig Medical Foundation (https://www.hartwigmedicalfoundation.nl/en/data/data-acces-request/). All the scripts and code is available on the Curtis Lab Github Repo: https://github.com/cancersysbio/germline-epitopes

## Authors’ contributions

**Project Initiation:** K.E.H., C.C.

**Bioinformatic Analyses:** K.E.H., A.K.

**Statistical Analyses:** K.E.H., N.F.G.

**Manuscript First Draft:** K.E.H.

**Supervised Research:** R.B.W., M.A., C.C.

**Manuscript Editing & Approval:** All authors

## Conflict of Interest Statement

Unrelated to this work, C.C. is an advisor to Genentech, Bristol Myers Squibb, 3T Biosciences, Resistance Bio, DeepCell, NanoString and has equity in DeepCell, Grail/Illumina.

## Acknowledgements

The authors thank members of the Curtis lab for helpful suggestions and support. The results described here are based in part upon data generated by the TCGA Research Network (http://cancergenome.nih.gov/) and the NCI Human Tumor Atlas Network. K.E.H was supported by a CIHR Banting Fellowship. Research in the Curtis Lab is supported by the NIH Director’s Pioneer Award (DP1-CA238296), a U54 Metastasis Network Center Grant (U54CA261719) and a Chan Zuckerberg Investigator award to C.C.

---

**Supplementary Figure 1.**
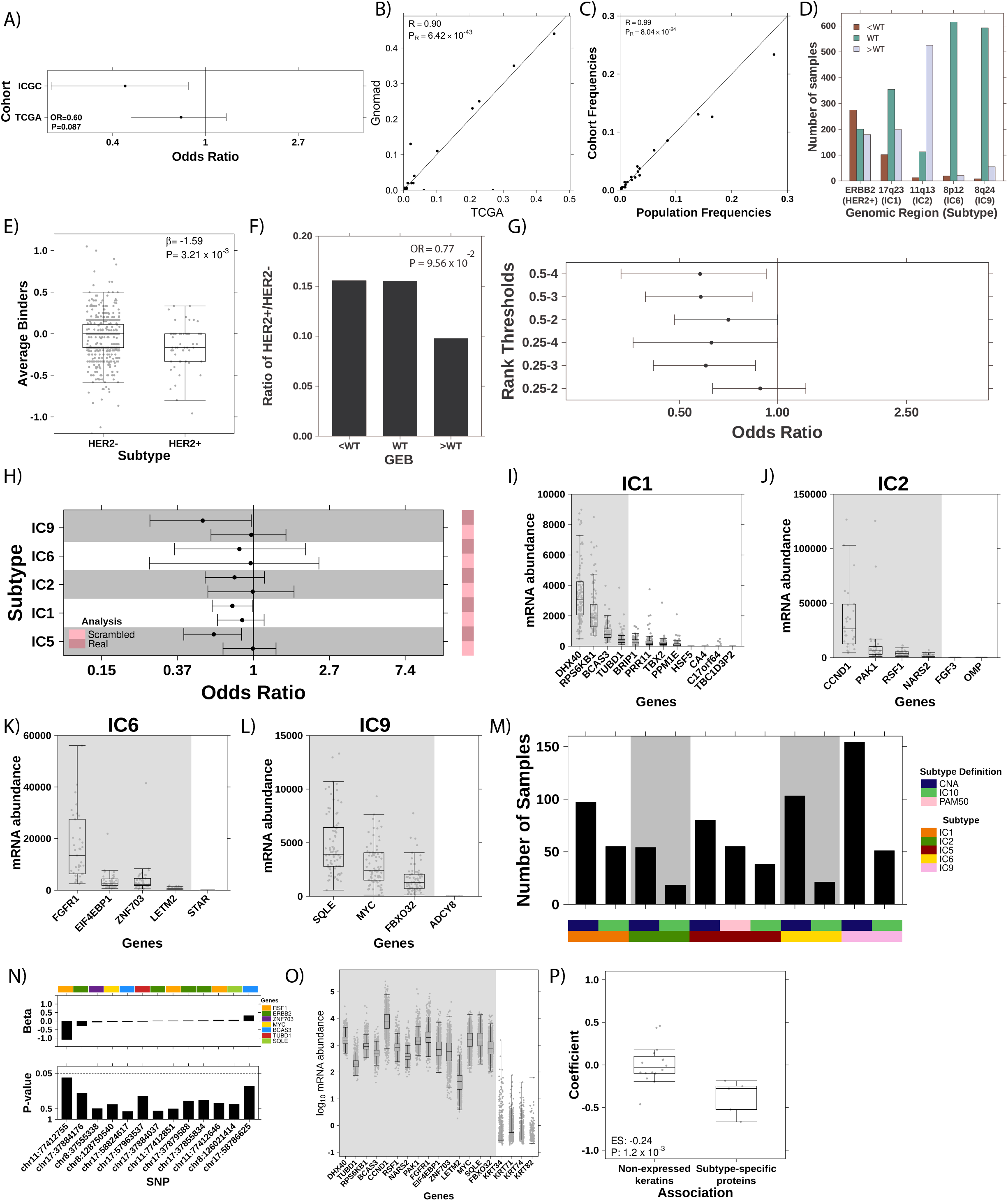
GEB in oncogene selects against oncogene amplification. **A)** Forest plot shows OR and 95% confidence intervals for association between number of HLA alleles an individual possesses that can bind GP2 and whether the individual has HER2+ breast cancer in ICGC and TCGA. **B)** Scatterplot of minor allele frequencies (MAF) in TCGA discovery cohort compared to population frequencies in Gnomad (Pearson correlation). **C)** Scatterplot of HLA allele frequencies in TCGA discovery cohort compared to population frequencies in The Allele Frequency Net Database. **D)** Barplot showing the number of samples that have low, medium or high GEB (defined as less than the reference genome (<WT), the same as reference (WT) or greater than the reference genome (>WT)) in subtype specific recurrently amplified loci. **E)** Boxplot showing depletion of the average number of binders in *ERBB2* in HER2+ breast cancer compared to HER2- breast cancer. Statistics from a logistic regression model correcting for the first six genetic principal components. Boxplot represents median, 0.25 and 0.75 quantiles with whickers at 1.5x interquartile range. **F)** Barplot shows the ratio of HER2+ to HER2- patients with low, medium or high GEB defining HER2+ as having an *ERBB2* amplification (*i.e.* >4 copies). Statistics from logistic regression model correcting for the first six genetic principal components. **G)** Scatterplot showing odds ratio (x-axis) between GEB and HER2+ breast cancer considering varying definition of HLA binders (y-axis). **H)** Negative association between GEB and subtype commitment is not driven by germline variants alone. Forest plot shows odds ratio and 95% confidence intervals for the true associations (“real”) compared to associations run with scrambled HLA alleles (“scrambled”). Odds ratio and 95% confidence interval plotted were calculated as the median, 0.025 and 0.975 quantiles of 1,000 iterations of scrambled HLA alleles. Covariate along the right indicates if statistics are from real or scrambled analyses. **I-L)** mRNA abundance of recurrently amplified genes in each of the four high risk ER+ IntClust subtypes: IC1 **(I)**, IC2 **(J)**, IC6 **(K)** and IC9 **(L)**. **M)** Boxplot shows the number of samples (y-axis) corresponding to each subtype (x-axis) based on alternative subtype definitions. **N)** Barplot shows effect size (top) and p-value (bottom) from association with breast cancer risk from Zhang et al. (*7*) **O-P)** As a negative control, we tested the association of the GEB in unexpressed keratins, KRT34, KRT71, KRT74 and KRT82, with the PAM50 subtypes. As these proteins are not expressed in mammary tissue, there should be no association. **(O)** Boxplot shows log10 mRNA abundance of subtype specific genes on the left (grey background shading) compared to the unexpressed keratins on the right. **(P)** Boxplot shows the coefficients for these analyses are significantly closer to zero compared to the coefficients from the subtype-specific protein analyses. Effect size (ES) represents the difference in medians. P-value from Mann Whitney Rank Sum Test.

**Supplementary Figure 2.**
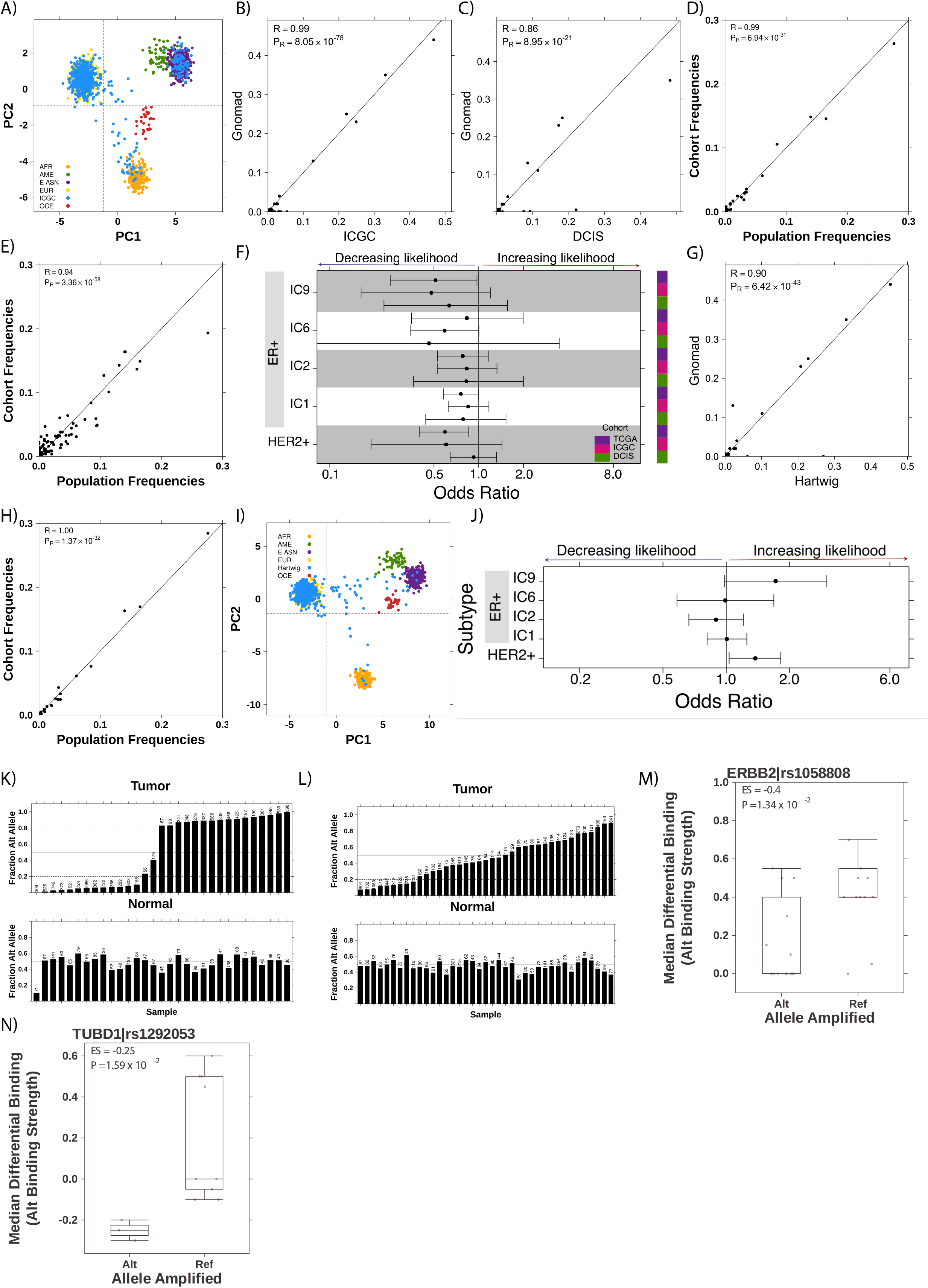
Germline-mediated immunoediting dictates breast cancer subtype early during tumorigenesis. **A)** Scatterplot shows principal component 1 and 2 for ICGC replication cohort against reference cohort based on 128 ancestry informative markers (*51*). Majority of samples cluster with European population. Dotted lines represent cutoffs for inclusion in analysis. **B-C)** Scatterplot of minor allele frequencies (MAF) in ICGC **(B)** and DCIS **(C)** replication cohorts compared to population frequencies in Gnomad (Pearson correlation). **D-E)** Scatterplot of HLA allele frequencies in ICGC **(D)** and DCIS **(E)** replication cohorts compared to population frequencies in The Allele Frequency Net Database. **F)** Across five subtypes and three independent cohorts, a high GEB in subtype-specific oncogenes is associated with decreased likelihood of developing the cognate subtype. Forest plot shows the odds ratio and 95% confidence intervals for three cohorts: DCIS, TCGA and ICGC. Covariate on the right indicates cohort. **G)** Scatterplot of minor allele frequencies (MAF) Hartwig cohort compared to population frequencies in Gnomad (Pearson correlation). **H)** Scatterplot of HLA allele frequencies in Hartwig cohort compared to population frequencies in The Allele Frequency Net Database. **I)** Scatterplot shows principal component 1 and 2 for Hartwig cohort against reference cohort based on 128 ancestry informative markers (*51*). Majority of samples cluster with European population. Dotted lines represent cutoffs for inclusion in analysis. **J)** Forest plot shows association between GEB and subtype in metastatic breast cancer (Hartwig). No association was observed in metastatic breast cancer. **K-L)** Barplots show fraction of reads supporting the alternative allele in the tumor (top) and the normal (bottom) for two common variants: rs1058808 **(K)** and rs1292053 (**L)**. The number on the top of each plot shows the top number of reads covering each loci. The horizontal line indicates fraction = 0.5 while the dotted lines represent fraction = 0.2 or 0.8. **M-N)** Boxplots of median differential binding per sample for epitopes derived from the alt allele *vs* ref allele (y-axis) for samples that preferentially amplified the alt or the ref allele. Effect size and p-value from Mann-Whitney rank sum test. Boxplots show analysis for rs1058808 derived from *ERBB2* **(M)** and rs1292053 derived from *TUBD1* **(N)**.

**Supplementary Figure 3.**
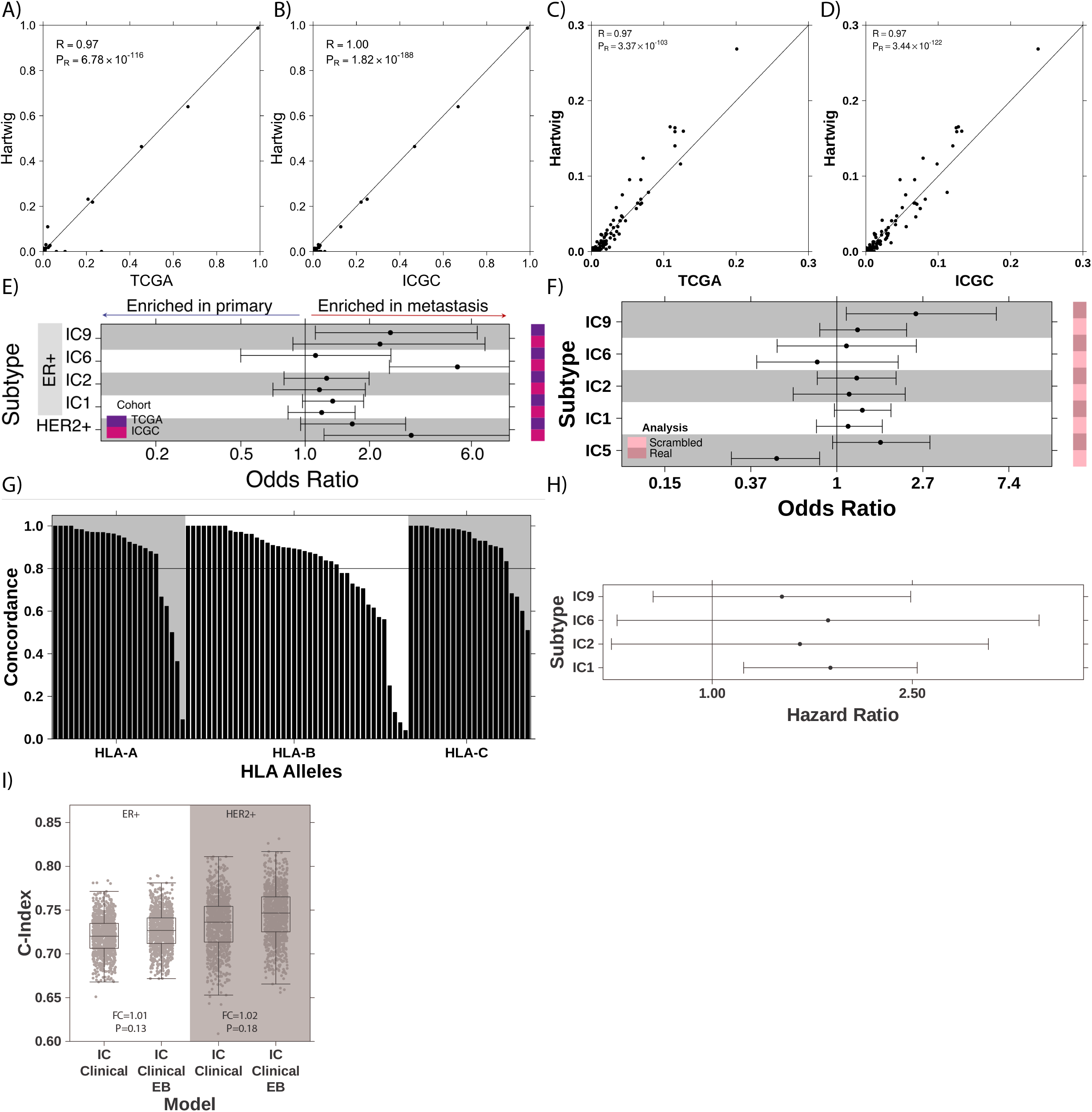
Tumors that overcome a high GEB are more aggressive A-B) Scatterplot of minor allele frequencies (MAF) in TCGA **(A)** and ICGC **(B)** compared to Hartwig (Pearson correlation). **C-D)** Scatterplot of HLA allele frequencies in TCGA **(C)** and ICGC **(D)** compared to Hartwig (Pearson correlation). **E)** Forest plot comparing GEB between primary and metastatic tumors of the same subtype. Two primary cohorts were evaluated, TCGA and ICGC, against one metastatic cohort (Hartwig). Covariate on the right indicate which cohort. **E)** Enrichment in metastatic tumors is not driven by germline variants alone. Forest plot shows odds ratio and 95% confidence intervals for the true associations (“real”) compared to associations run with scrambled HLA alleles (“scrambled”). Odds ratio and 95% confidence interval plotted were calculated as the median, 0.025 and 0.975 quantiles of 1,000 iterations of scrambled HLA alleles. Covariate along the right indicates if statistics are from real or scrambled analyses. **G)** Accuracy of HLA imputation from TCGA SNP6 data compared to HLA genotyping from WES from Polysolver (*49*) as the gold standard. Horizontal line indicates accuracy of 80%. **H)** Forest plot shows hazard ratio (HR) and 95% confidence intervals from CoxPH correcting for the first two genetic principal components, age and percent genome altered (PGA) for each high-risk ER+ subtype individually. **I)** Boxplot shows c-index of predictive models considering Integrative Clusters (IC) and clinicopathologic features (age, size, grade and node involvement) alone or in combination with GEB for 1,000 bootstrapped iterations. Fold change (FC) is calculated as the ratio of medians while the p-value is calculated as 1 – the proportion of iterations where the c-index of the IC+clinical+GEB model was greater than the IC+clinical model.

**Supplementary Figure 4.**
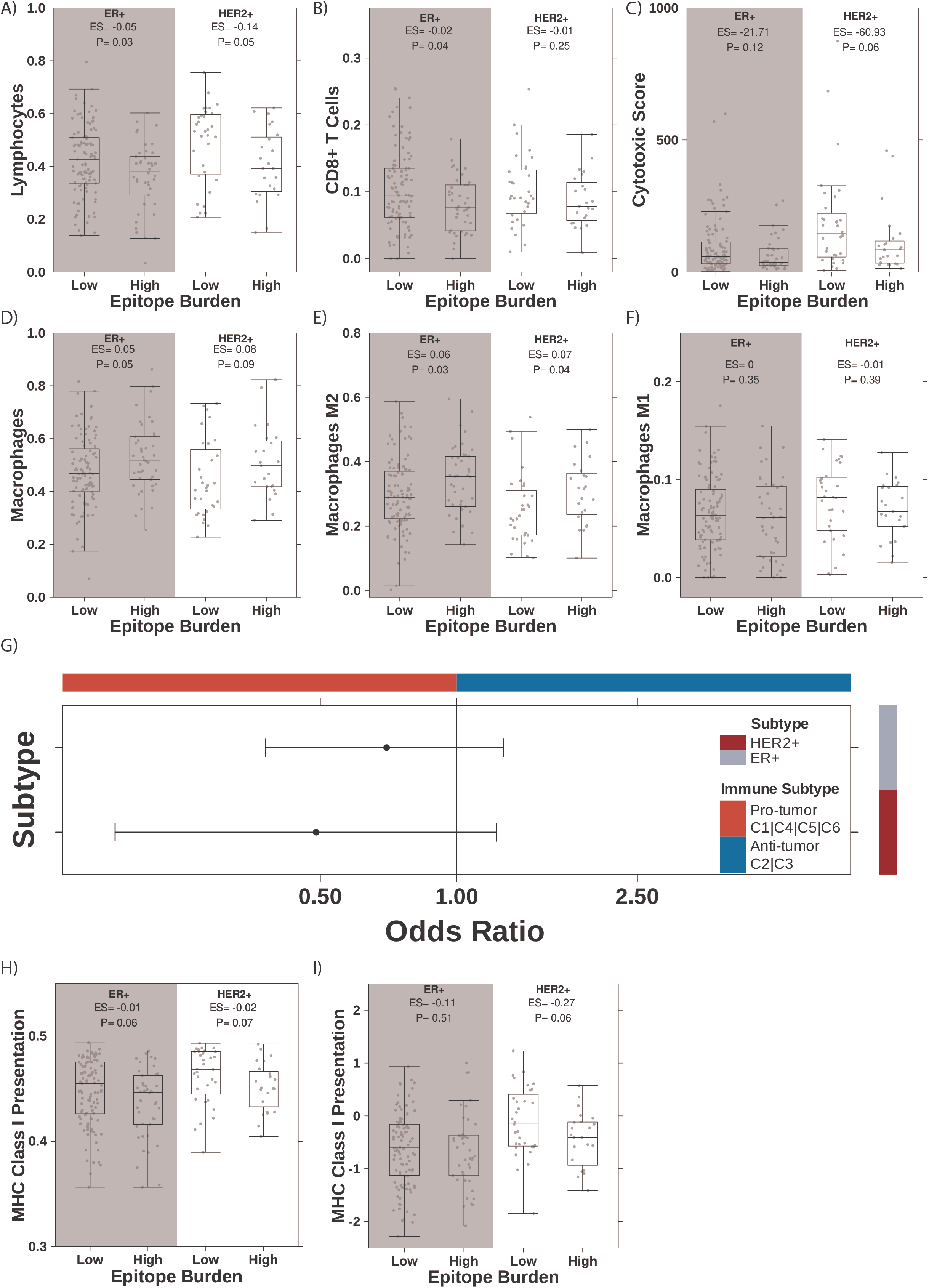
**A high GEB promotes an immunosuppressive phenotype A-F)** Lymphocyte infiltration **(A)**, CD8+ T cells infiltration **(B)**, cytotoxic score **(C)**, macrophage infiltration **(D)**, M2- **(E)** or M1-polarized macrophages **(F)** in ER+ or HER2+ high germline epitope tumors compared to low germline epitope tumors in TCGA. Effect size (ES) shows difference in medians while p-value is from Mann-Whitney Rank Sum test. Boxplot represents median, 0.25 and 0.75 quantiles with whickers at 1.5x interquartile range. **g)** Forest plot shows odds of developing anti-tumor immune subtype (x-axis) given a high GEB in HER2+ or ER+ subtypes (y-axis). Covariate along the right indicate the subtype evaluated while the covariate along the top indicates the interpretation of the direction of effect. **H-I)** Boxplot of MHC Class I antigen presentation pathway measured by two different transcriptional signatures (y-axis) stratified by high *vs* low GEB tumors (x-axis). Effect size (ES) and p-value from Mann-Whitney Rank Sum test.

**Supplementary Table 1 GEB across five individual breast cancer cohorts** Patient-level GEB in five recurrent amplicons for four individual cohorts: TCGA (primary invasive breast cancer), ICGC (primary invasive breast cancer), DCIS, Hartwig (metastatic breast cancer) and METABRIC (primary invasive breast cancer). Table includes subtype annotations, GEB in each of the four amplicons, genetic principal components and number of somatic SNVs for each cohort. METABRIC table additionally includes overall survival at five years and percent genome altered (PGA).

**Supplementary Table 2 Immune landscape of high *vs* low GEB tumors** Immune transcriptomic features from Thorsson et al. (*42*) for HER2+ and ER+ high *vs* low GEB tumors.

